# Complementation testing identifies causal genes at quantitative trait loci underlying fear related behavior

**DOI:** 10.1101/2024.01.03.574060

**Authors:** Patrick B. Chen, Rachel Chen, Nathan LaPierre, Zeyuan Chen, Joel Mefford, Emilie Marcus, Matthew G. Heffel, Daniela C. Soto, Jason Ernst, Chongyuan Luo, Jonathan Flint

## Abstract

Knowing the genes involved in quantitative traits provides a critical entry point to understanding the biological bases of behavior, but there are very few examples where the pathway from genetic locus to behavioral change is known. Here we address a key step towards that goal by deploying a test that directly queries whether a gene mediates the effect of a quantitative trait locus (QTL). To explore the role of specific genes in fear behavior, we mapped three fear-related traits, tested fourteen genes at six QTLs, and identified six genes. Four genes, *Lsamp, Ptprd, Nptx2* and *Sh3gl,* have known roles in synapse function; the fifth gene, *Psip1,* is a transcriptional co-activator not previously implicated in behavior; the sixth is a long non-coding RNA *4933413L06Rik* with no known function. Single nucleus transcriptomic and epigenetic analyses implicated excitatory neurons as likely mediating the genetic effects. Surprisingly, variation in transcriptome and epigenetic modalities between inbred strains occurred preferentially in excitatory neurons, suggesting that genetic variation is more permissible in excitatory than inhibitory neuronal circuits. Our results open a bottleneck in using genetic mapping of QTLs to find novel biology underlying behavior and prompt a reconsideration of expected relationships between genetic and functional variation.

## INTRODUCTION

A major challenge in behavior genetics is to turn genetic information into mechanistic understanding, of the sort that would for example be useful in designing new treatments for psychiatric disorders and more generally in understanding how genetic variation leads to behavioral variation. A key step is progressing from quantitative trait locus (QTL) to gene, which has only been achieved for any complex trait, in any species, in a small number of cases^1,2^. Of the 5,000 QTLs identified in rodents, less than 100 genes have been identified, almost all on the basis of correlative evidence, such as proximity of a gene to the QTL or alterations in transcript abundance, rather than by a causal test of a gene’s candidacy^2^. Currently, there is no consensus on how to proceed from quantitative trait locus (QTL) to gene.

Here we demonstrate the power of a quantitative complementation (QC) test, first applied in Drosophila^3^ and later shown to work in rodents^4^, to directly query the causal gene impacted by the QTL. Construction and phenotyping of F1 hybrids with and without a knockout of a candidate gene, and of inbred strains with and without the knockout of a candidate gene, test whether the QTL operates through the gene under investigation. By separately assaying the joint effect of QTLs (from the phenotypic difference between strains) and the effect of the mutation (from the phenotypic difference between knockouts and wildtype), the QC test reveals a QTL whose effect depends on the presence of the candidate gene as a significant interaction between the effect of mutation and the effect of strain. Applying the test requires access to inbred animals carrying a knockout on the same genetic background as the QTL, which has been difficult to achieve when knockouts were generated using homologous recombination in 129 strains or C57BL/6N. The development of the CRISPR-Cas9 technology now lifts that restriction by enabling genetic manipulation in any strain^5,6^.

We set out to apply the QC test to fear-related behaviors. We chose these traits because we could use extensive information available about the brain regions involved (ventral hippocampus and amygdala^7,8^), and underlying circuitry^9–12^, to explore where, and how in the brain genetic variation results in behavioral change. We reasoned that by knowing the causal genes, we could identify changes in their expression and regulation at a single cell level, revealing the molecular mechanisms that mediate the impact of genetic variants on traits.

## RESULTS

### Generating a large-scale set of phenotypes and genotypes

Our first task was to identify QTLs involved in fear-related behaviors that ideally indicated one or a small number of genes for QC testing. We chose a hybrid mouse diversity panel (HMDP) for mapping, because of its potential to deliver high resolution^2^ (thus increasing the chance of finding QTLs containing a small number of candidate genes), and because it includes recombinant inbreds derived from C57BL/6J, hence providing many QTL alleles on a strain readily amenable to CRISPR-Cas9 modification (the HMDP strains and numbers of animals used are listed in Table S1). We mapped variation in a conditioned fear assay in which animals were exposed to an auditory cue associated with an aversive shock. Re-exposure to the auditory cue elicits a fear response, measured as the amount of time spent freezing ^13,14^ and referred to in this paper as FC-cue. The context in which conditioning occurs also elicits freezing (FC-context)^15^, and we assayed this too. We also mapped variation in an unconditioned fear assay using the elevated plus maze (EPM), whose open arms act as anxiogenic environment. We counted the number of entries into the EPM’s open arms as a measure of fear, referred to henceforth as EPM-open.

To enable us to perform a well-powered meta-analysis and robustly identify loci at high resolution we combined our data with those from 25 publications^16–40^, four unpublished data sets at the Jackson laboratory website and one unpublished data set at the Gene Network website (summarized in Table S2). We implemented a rigorous pipeline to harmonize phenotypes across the multiple data sets (Supplemental Material), generating phenotypes on a total of 6,544 mice from seven cohorts (Table S2), that to our knowledge comprise the largest data set assembled for genetic mapping of fear-related behavior.

Our next requirement was a set of genotypes at markers sufficiently dense to capture most causative variants. Short read sequencing provides almost complete catalogs of genetic variants for sixteen inbred strains^41^, which are also a resource for imputing variants in the other mapping strains^42^. We carried out imputation using a hidden Markov model based technique with proven effectiveness in inbred model organisms^43^. After imputation, we obtained genotypes on 16,767,664 markers (the imputation strategy and validation results are described in Supplemental Material). For mapping, marker sets were chosen that are polymorphic within the strains of each data set, a number varying from 4 to 16 million single nucleotide polymorphisms (SNPs) segregating between the inbred strains, sufficient to identify causative variants at QTLs^44,45^.

### Genetic mapping of conditioned and unconditioned fear

The phenotypic dataset we obtained is highly structured, consisting of closely related individuals (recombinant inbreds share half their genome, like full siblings) as well as distantly related individuals. Genome wide association was carried out with GEMMA, which implements a linear mixed model marker association test ^46,47^, including a genomic relationship matrix to control for the population structure arising from the inclusion of individuals with different degrees of relatedness. We mapped each phenotype in each cohort and carried out a meta-analysis of the results using a random effects model implemented in METASOFT ^48,49^. We used Genome Reference Consortium Mouse Build 38 (GCA_000001635.2, mm10), and all coordinates in this paper refer to that build.

To assess significance we used GCTA ^50^ to simulate data for each cohort, and meta-analyzed the results in the same way as we did for the real data sets. From 100,000 simulations we obtained a 5% threshold of negative logarithm (base 10) of the association P-value (logP) of 4.50, consistent with thresholds estimated by others using the same experimental design ^32,51^. Applying this significance threshold we identified 14 loci for FC-context, 15 for FC-cue and 72 for EPM-open (Table S3). At a threshold corrected for testing three phenotypes (logP = 4.97) the number of loci for EPM-open fell to 60, while remaining unchanged for the other two phenotypes. Four loci were common to FC-context and FC-cue and four common to EPM-open and FC-cue, giving 93 unique loci. The Manhattan plots in Figure 1 show all loci that exceeded the logP threshold of 4.5.

**Figure 1:**
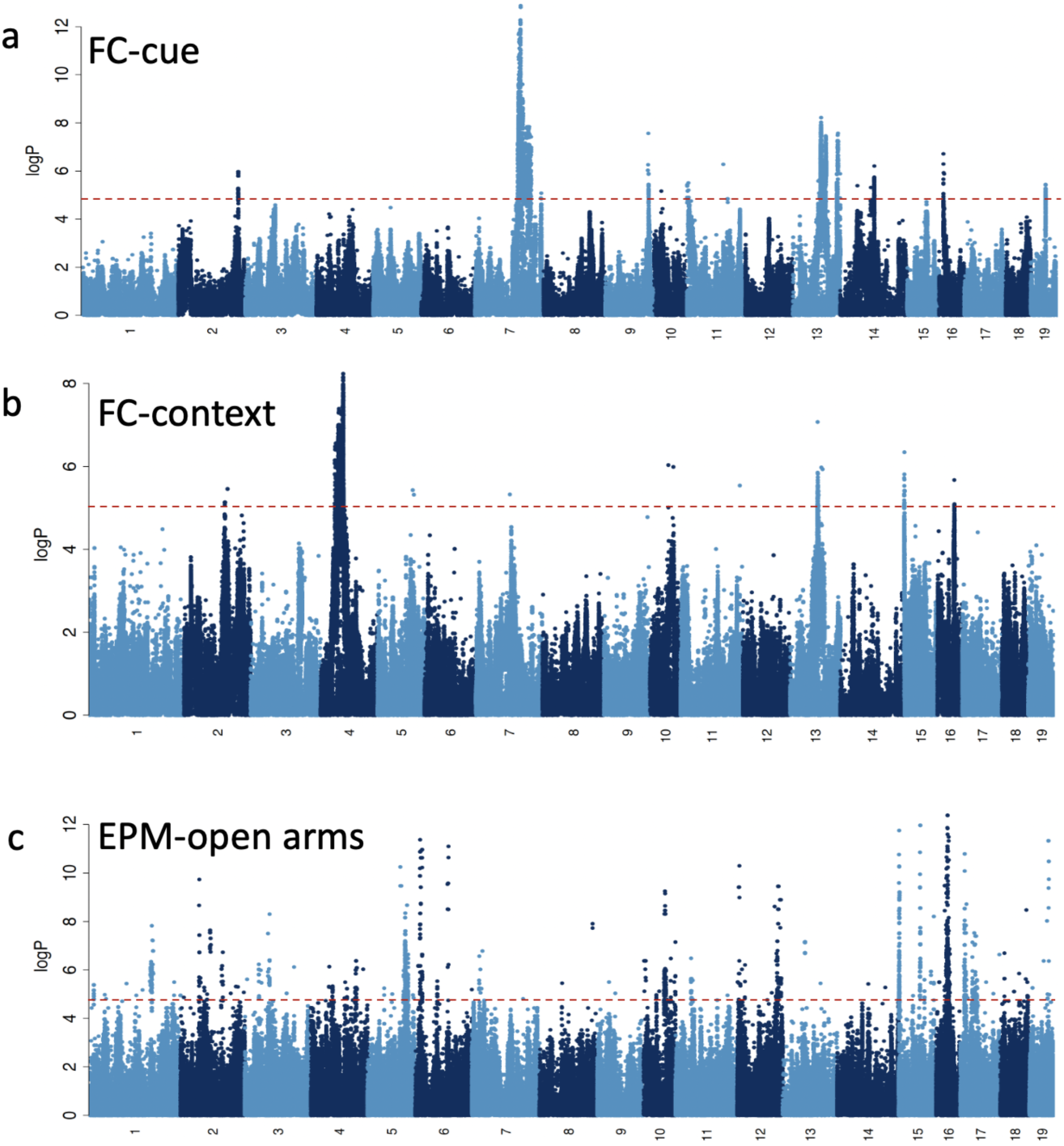
Manhattan plots for three fear-related behavior in mice. The data show the results of a meta-analysis of a) 1,931 mice for cue conditioning (FC-cue), b) 2,671 for contextual conditioning (FC-context) and c) 1,942 for entries into the open arms of the elevated plus maze (EPM-open arms). Chromosome numbers are listed on the horizontal axis. The vertical scale is the negative logarithm (base 10) of the association P-value. A horizontal dotted red line gives the location of the 5% significance threshold (from simulation).

While the genetic architecture of fear related behaviors was highly polygenic, we found evidence for a few large effect loci, notably on chromosomes 7 and 13 for FC-cue (Figure 1a) and chromosomes 4 and 13 for FC-context (Figure 1b). More than five times as many QTLs were identified underlying EPM-open than either of the fear-conditioning phenotypes, likely reflecting higher polygenicity of the trait (Figure 1c). The size of the QTL intervals varied considerably, from 0.02 to 19.6 megabases (Mb) (confidence intervals estimated by simulation^52^) with a median of 2.5 Mb, each locus containing a median of 23 genes (range from 0 to 329, Table S3).

### QC testing identifies six genes for fear related behavior

Which of the genes at a locus is causal for the phenotype? To answer this for specific loci, we first chose a 2 Mb QTL at the end of chromosome 13 associated with FC-cue (logP 7.6), previously identified in a panel of BXD recombinant inbreds^26^, that includes a hyperpolarization-activated cyclic nucleotide-gated channel 1 (*Hcn1*). Since pharmacological blockade of HCN1 reduces freezing, *Hcn1* has been proposed as the causal gene at this locus^26^ (although deletion of *Hcn1* does not alter conditioned fear ^53^). We used an exon-excision CRISPR-Cas9 strategy to knockout out all four annotated and one unannotated gene lying within the 95% confidence intervals of the QTL (Figure 2a) (we omitted the single exon unannotated transcript *Gm6416*). We used a quantitative polymerase chain reaction to confirm that the engineered mutations altered RNA abundance. The CRISPR-Cas9 strategy and characterization of the mutants is described in Supplemental Material.

**Figure 2.**
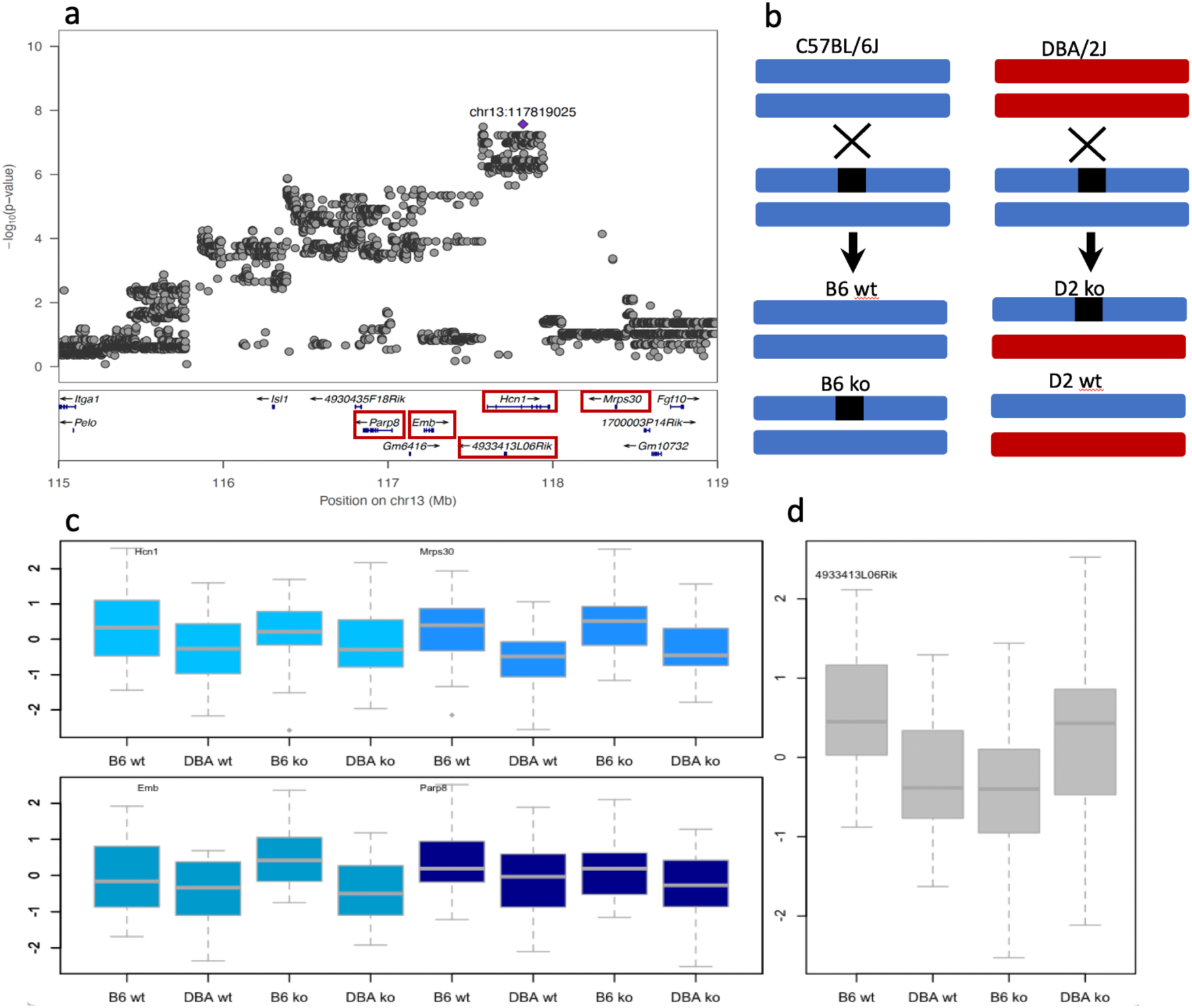
Quantitative complementation of five genes at a locus on chromosome 13 for cue fear conditioning. Panel a) provides QTL regional information from LocusZoom ^54^. The top part of the panel shows the association results from a meta-analysis with the position of the highest scoring variant annotated in purple. The vertical scale is the negative logarithm (base 10) of the association P-value. The bottom section of panel a) gives the location and orientation of genes at the locus. A red box identifies each gene used in the quantitative complementation tests. b) Design of the quantitative complementation test. Black boxes indicate the knockout allele and strain is indicated by color, where blue is C57BL/6J and red is DBA/2J. An ‘X’ indicates a cross between the named groups above and below the ‘X’, and an arrow points to the progeny of each cross. The four groups used in the quantitative complementation test are wild type C57BL/6J (B6 wt), heterozygote knockouts (B6 ko), an F1 from crossing DBA/2J to C57BL/6J (D2 wt), and the F1 knockout by DBA/2J (D2 ko). c) Results of quantitative complementation testing of four annotated genes in the region. Groups are B6.wt: wild type C57BL/6J; B6.ko: heterozygote knockouts on C57BL/6J; D2.wt: F1 from crossing DBA/2J to C57BL/6J; D2.ko: the F1 from crossing the knockout onto DBA/2J. These genes were not significant by quantitative complementation (Table 1). d) Results of the quantitative complementation test for the long non-coding RNA *4933413L06Rik*. The difference between these groups yielded a significant interaction result in the quantitative complementation test (Table 1).

Each knockout was subject to a QC test (Figure 2b), to find out which gene was mediating the effect of the locus on FC-cue. The results are shown in Figure 2. Four genes tested negative (Figure 2c), while one was positive (Figure 2d), which to our surprise was an unannotated long non-coding RNA, *4933413L06Rik*. To interpret panels c and d in Figure 2, consider that the first two groups in Figure 2b (B6.wt and DBA.wt) measure a strain effect, arising from a single copy of the D2 genome in the F1 animals (assuming strict additivity, the strain effect should be about half that found in inbred strain comparisons, consistent with our results). Any difference between the B6.wt and B6.ko groups is attributable to the presence of the knockout.

The pattern of results in Figure 2c for the four genes tested looks identical: a strain effect can be seen but no discernable difference between groups carrying the knockout and the wildtypes. By contrast, Figure 2d shows a different pattern. Animals with the knockout do differ from their respective wild types, indicating that the knockout has an effect on the phenotype, however with differences between the two strains. The B6.ko animals spend less time freezing than their wild type siblings (B6.wt), while the DBA.ko group freeze *more* than their wild types (DBA.wt). In other words, the effect of the mutation depends on the strain background, which we assume to be due to a nearby QTL.

We tested this relationship between strain and knockout by analysis of variance and found a significant interaction (P = 0.0015, Table 1), indicating that the effect of the QTL is mediated by the unannotated gene *4933413L06Rik*. We found no evidence for the involvement of *Hcn1*, demonstrating that QC testing can unambiguously identify causal genes at QTL.

**TABLE 1:**
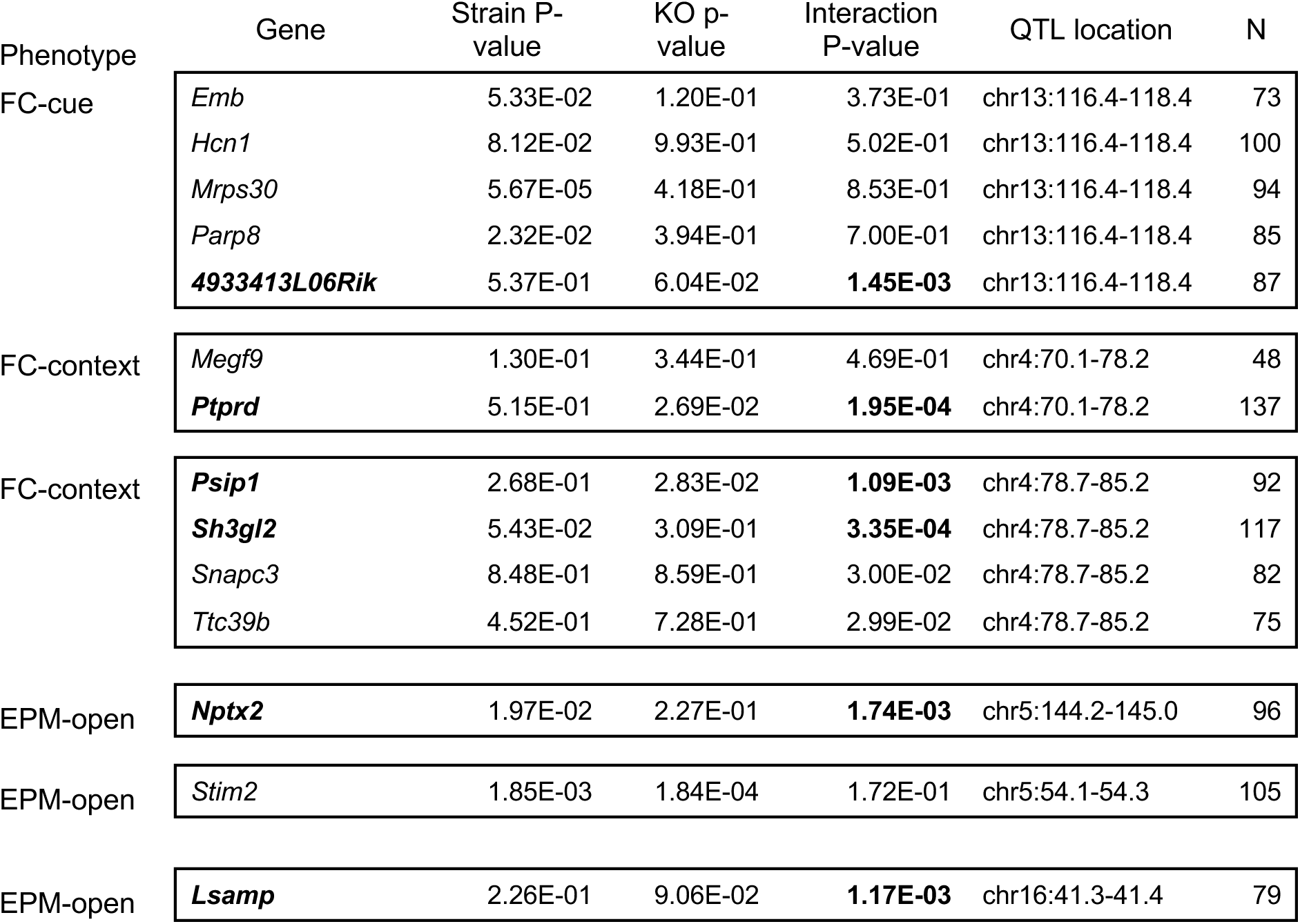
Quantitative complementation testing of fourteen genes at six QTLs for fear-related behavior. The table shows the P-values of an analysis of variance testing for an interaction between strain and knockout (KO) in the four groups of the quantitative complementation test. QTL locations are given in Mb and the name of the phenotype tested. Results exceeding a multiple testing corrected P-value of 0.0035 are shown in bold. N is the total number of animals used for the quantitative complementation test. QTL location coordinates are to mouse Genome Reference Consortium Mouse Build 38 (mm10).

We next tackled the most significant region of association for FC-context, a 15 Mb region on chromosome 4 (between 70 and 85). There were two QTLs here (70-78Mb and 79-85Mb, Figure 3a and 3c), together containing 46 genes. To narrow the choice of genes, we tested genes close to the most highly associated markers and with confirmed gene expression in the hippocampus and amygdala (from published sources^55–62^). At the first locus, the interaction P-value of *Ptprd* exceeded a multiple correction testing threshold (P < 0.0036 for testing 14 genes listed in Table 1) while at the second two genes, *Psip1* and *Sh3gl3*, exceeded the threshold (results shown in Figure 3b and 3d, and Table 1).

**Figure 3.**
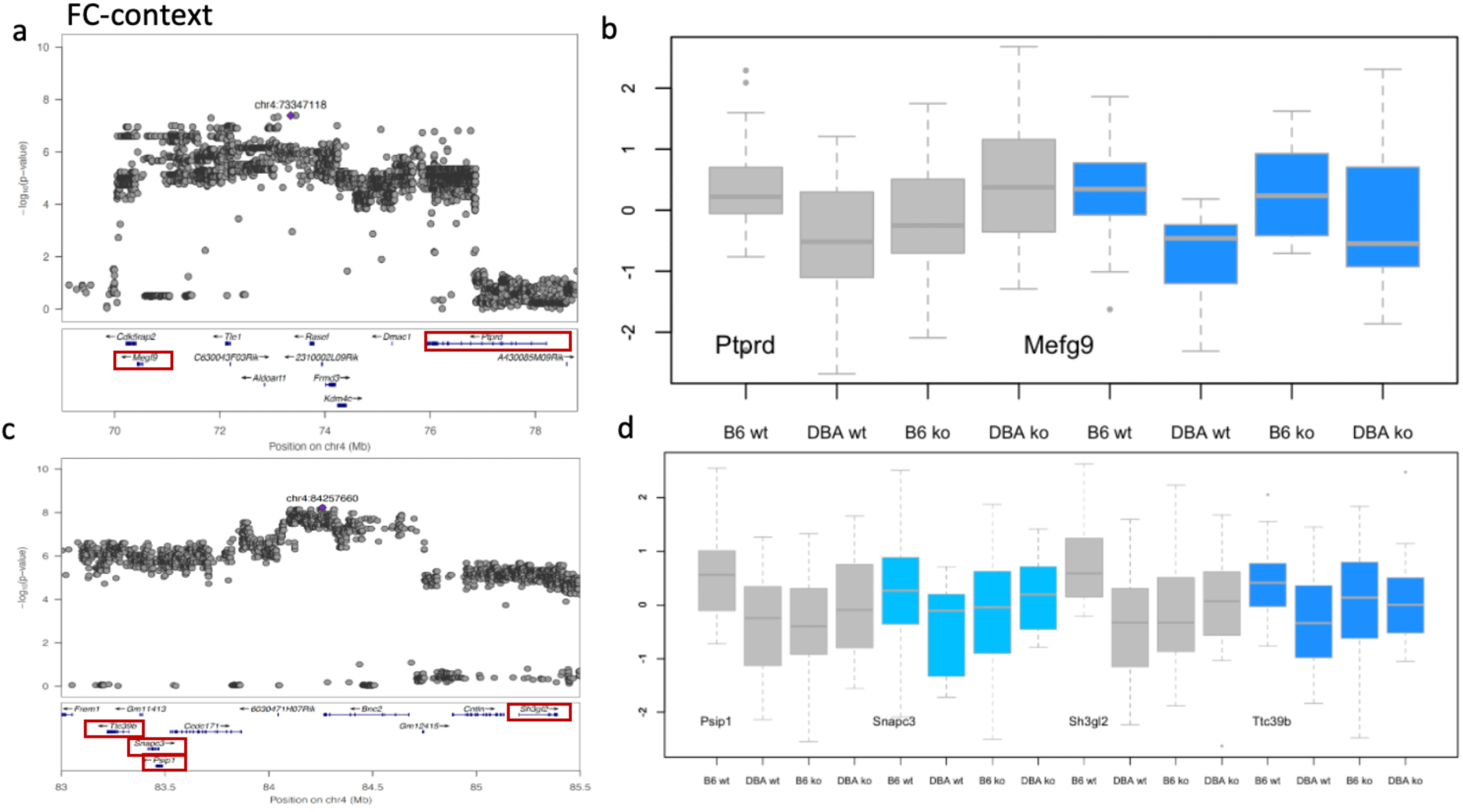
Quantitative complementation of four genes at two QTLs on chromosome 4 for contextual fear conditioning. Panels a) and c) display QTL regional information from LocusZoom^54^. The top part of the panel shows the association results from a meta-analysis with the position of the highest scoring variant annotated in purple. The vertical scale is the negative logarithm (base 10) of the association P-value. The bottom section of the panel section gives the location and orientation of genes at the locus. A red box identifies each gene used in the quantitative complementation tests. Panels b) and d) show results of quantitative complementation testing of six annotated genes. Groups are B6.wt: wild type C57BL/6J; B6.ko: heterozygote knockouts on C57BL/6J; D2.wt: F1 from crossing DBA/2J to C57BL/6J; D2.ko: the F1 from crossing the knockout onto DBA/2J. Genes significant by quantitative complementation are shown in grey (results in Table 1).

Finally, we chose three loci contributing to variation in EPM-open behavior (Figure 4). The first, on chromosome 16, had a highly significant association (logP 9.48), and lies in an intron of a cell adhesion molecule, *Lsamp* (limbic system-associated membrane protein), whose deletion is known to alter behavior in the elevated plus maze ^63,64^. We found a significant interaction (P = 0.001, Table 1), indicating that the effect of the QTL is mediated by the *Lsamp* gene. We tested *Stim2* and *Nptx2* at two other loci, using the same CRISPR-Cas9 exon deletion approach. While the QC test confirmed the candidacy of *Nptx2,* it was negative for *Stim2* (Table 1).

**Figure 4.**
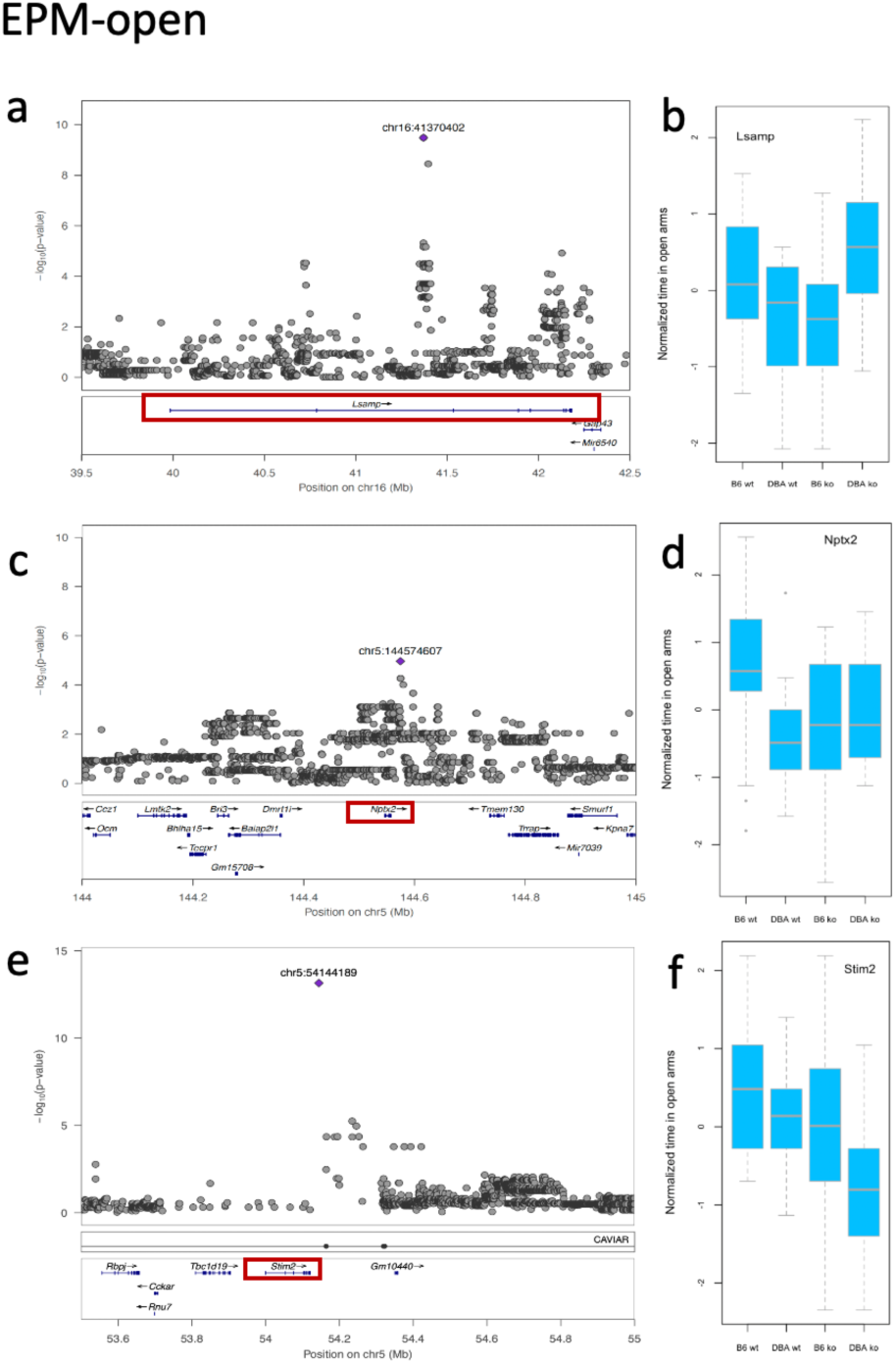
Quantitative complementation of three genes at three QTLs for entries into the open arms of the elevated plus maze. contextual fear conditioning. Panels a), c) and e) display QTL regional information from LocusZoom^54^. The top part of the panel shows the association results from a meta-analysis with the position of the highest scoring variant annotated in purple. The vertical scale is the negative logarithm (base 10) of the association P-value. The bottom section of the panel section gives the location and orientation of genes at the locus. A red box identifies each gene used in the quantitative complementation tests. Panels b), d) and f) show results of quantitative complementation testing. Groups are B6.wt: wild type C57BL/6J; B6.ko: heterozygote knockouts on C57BL/6J; D2.wt: F1 from crossing DBA/2J to C57BL/6J; D2.ko: the F1 from crossing the knockout onto DBA/2J.

Table one summarizes the results of the QC tests for all genes at the six loci we examined.

### Strain differences in gene expression and epigenetic regulation of causal genes is concentrated in excitatory hippocampal neurons

One hypothesis about how QTLs act is that they alter gene expression, which in our case would manifest as a difference in the abundance of transcripts from C57BL/6J and DBA/2J genomes in relevant cell types. With six genes in hand, we set out to test this assumption. We generated single-nucleus (sn) RNA-seq datasets in C57BL/6J and DBA/2J animals from the ventral hippocampus and amygdala. We identified 17 cell types in ventral hippocampus and 22 in amygdala with two replicates per strain per region (Figure 5, joint uniform manifold approximation and projection (UMAP)). The clusters included excitatory (glutamatergic) and inhibitory (GABAergic) neurons, as well as several types of non-neuronal cells that were annotated based on known marker genes (Methods). We define individual neuronal cell *types* based on neurotransmitter expression and region or top marker gene (for example Exc-DG in ventral hippocampus and Exc-Tafa1 in amygdala), and we define neuronal cell *class* as either excitatory or inhibitory through combining all excitatory or all inhibitory cell types respectively (one amygdala neuronal type was excluded from either cell class due to expression of both excitatory and inhibitory markers). Non-neuronal cell types were not further defined due to an enrichment step for neuronal nuclei during sample processing limiting the number of non-neuronal nuclei captured.

**Figure 5.**
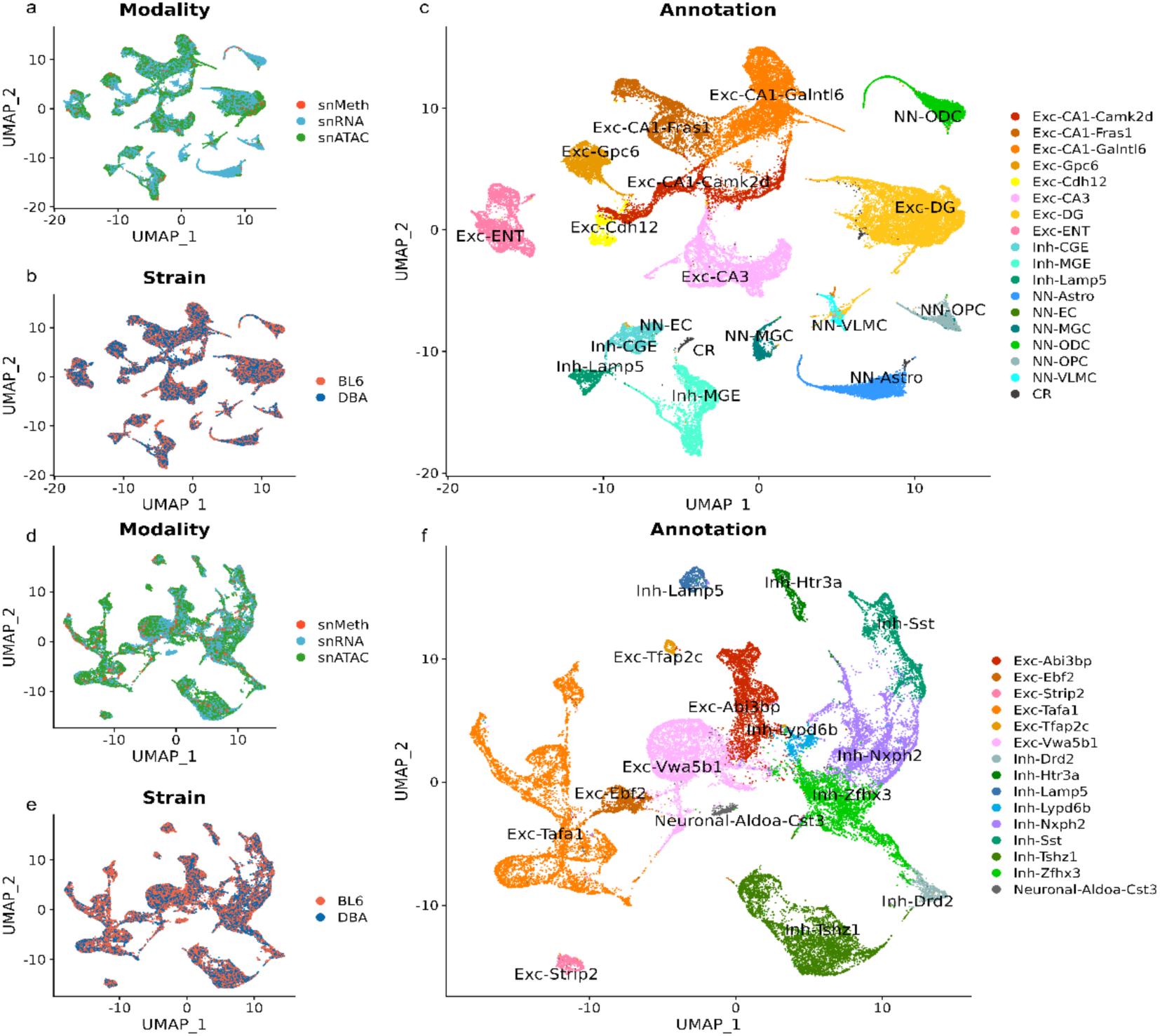
Cell type UMAP for ventral hippocampus and amygdala in B6 and DBA from single-nucleus RNA, ATAC, and methylation sequencing. Ventral hippocampal clusters plotted by a) modality, b) strain, and c) cell type identity. Amygdala neuronal clusters plotted by d) modality, e) strain, and f) cell type identity.

Given that QTL action is believed to have its effect on transcription through alteration of sequence at regulatory regions, we also generated single nucleus data for CpG methylation ^65^ and from a single nucleus Assay of Transposase Accessible Chromatin (snATAC-seq) ^66^. Both epigenetic modalities can be used to identify regulatory elements, such as promoters and enhancers^67^. Cell type clusters generated from single nucleus RNA profiles were propagated to snATAC and methylation datasets through joint embedding (Figure 5). We confirmed that cell types were integrated by modality (Fig 5a, d), that there was no obvious strain bias for cell types (Fig 5b, e), and that cell types were well-defined (Fig 5c, f; see Methods). Importantly for interpreting our analyses, the three epigenetic data sets were obtained independently from different animals, with two independent replicates per modality.

Five out of six genes showed significant differences in expression between C57BL/6J and DBA/2J (significance defined as exceeding a 5% P-value (logP > 1.3), adjusted for the total number of transcripts tested, from DEseq2^68^ output) : *4933413L06Rik* (in eleven cell types), *Lsamp* (in eleven cell types), *Psip1* (in five cell types), Nptx2 (in one cell type), and *Ptprd* (in eight cell types). Of the 36 cell types exhibiting differential expression for these genes, 31 were within the hippocampus. Different hippocampal cell types showed different degrees of overlap for differential gene expression of genes; for example, four of the five genes were differentially expressed within Exc-CA1-Galntl6 cells while only *4933413L06Rik* was differentially expressed within Exc-CA1-Camk2d cells. We noted that 22 of the 31 significant differences in the hippocampus and 3 out of 5 significant differences in the amygdala occurred in cell types categorized as excitatory neurons. Results are given in Supp Table S4.

We asked if the pattern of gene expression also held for snATAC-seq and methylation profiles, namely showing more strain differences in the hippocampus with an enrichment in excitatory neurons. To do this we made an inventory of variable snATAC-seq and methylation sites. We counted differential ATAC sites (again using a 5% threshold from adjusted P-values derived from DESeq2 ^68^) that contained sequence variants, on the assumption that differences at these sites were more likely genetic in origin than those without such differences (more than 90% of the significantly different ATAC sites contain a sequence variant, compared to 35% of non-variable sites). For the methylation data, we counted the number of fully methylated CpG sites in each cell type and divided them into those with and without mutations that disrupted methylation.

Figure 6 plots differential methylation and ATAC sites in the hippocampus and amygdala at the five QTLs where we identified causal genes. Three general observations arose from the data presented in Figure 6. First, there were more strain differences in the hippocampus than amygdala, as reflected in the greater number of dots compared to crosses. Second, at each locus the strain differences were found primarily in excitatory neurons, in both hippocampal and amygdala, as shown by the preponderance of red. Third, the pattern of strain differences for each modality varied between loci and genes, providing no unequivocal location or molecular signature to identify causal variants. The *Ptprd* locus (Fig 6a) contained a concentration of methylation, ATAC sites and sequence differences at the 3’ end of the gene, suggesting a location for one or more causal variants here. At the *Psip1* and *Sh3gl2* locus (Figure 6b), strain differences were spread across the region, with no indication of which might be relevant. The QTLs containing *4933413L06Rik* (Figure 6c) and *Nptx2* (Figure 6d) included a concentration of strain differences in methylation sites around the genes but with no obvious candidates for causative variants. One possible exception was the single ATAC site that exceeded a corrected significance threshold (logP 6) in an intron of the *Lsamp* gene (Figure 6e) coinciding with a 2kb deletion in DBA/2J. The function of this site, and the consequences of its deletion, are not known.

**Figure 6.**
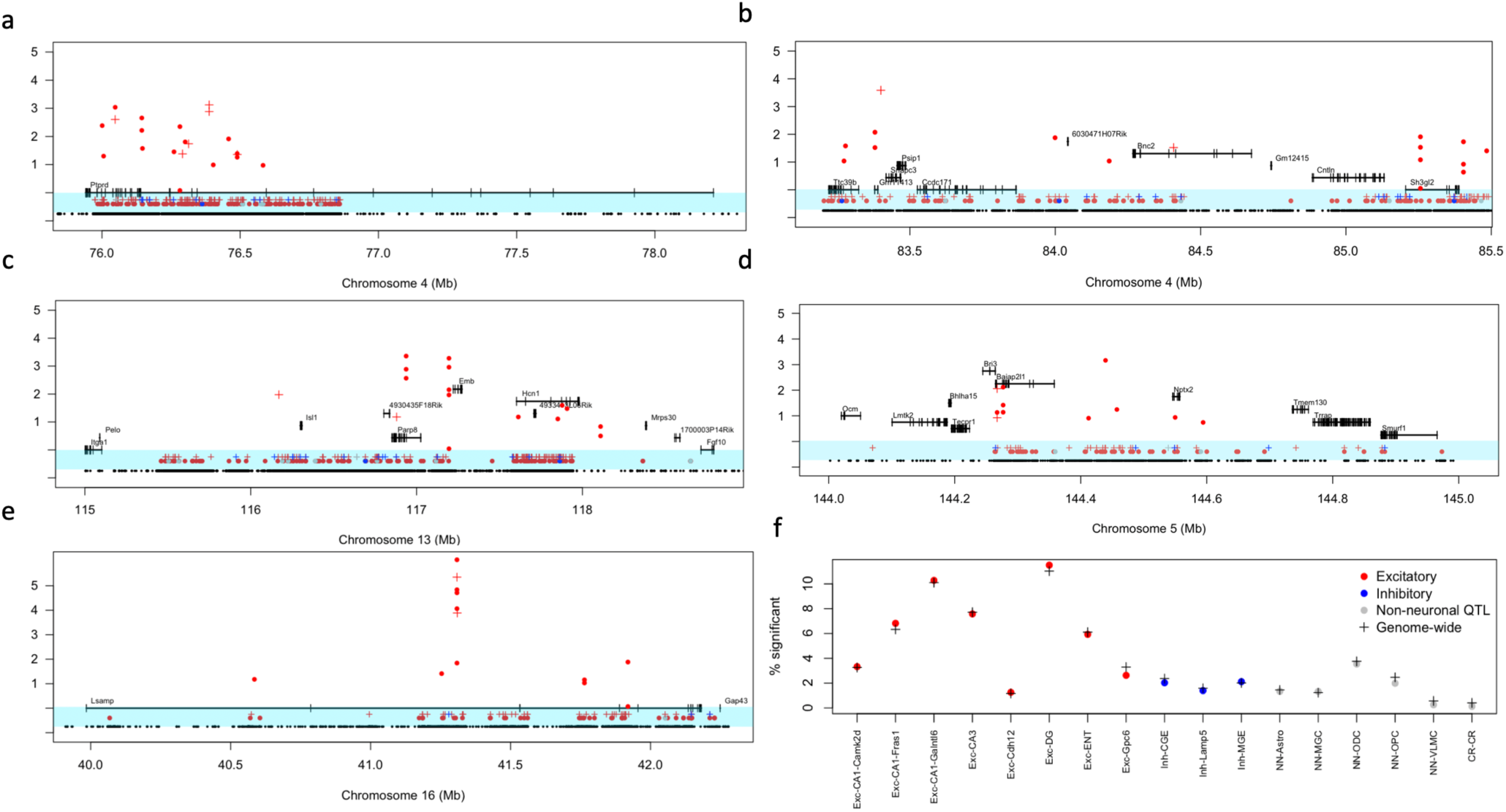
ATAC seq and methylation data at five QTLs in the ventral hippocampus and amygdala. In panels a) to e) data from the hippocampus is marked by a dot ( • ) and data from the amygdala by a cross ( + ). In each panel the lowest row shows the distribution of genetic variants (black). The next two tracks, immediately below the genes and indicated by the blue background, show the location of methylation sites that coincide with a sequence variant and are thus present in only one of the two strains. These sites are colored according to the class of cell type in which they occur: red for excitatory neurons, blue for inhibitory neurons and grey for non-neuronal cell types. Above the methylation tracks are shown ATAC sites which differ significantly between the strains. The vertical axis shows the absolute value of log2 fold change for these ATAC sites. Sites are again colored according to the class of cell type in which they occur, using the same color scheme as for methylation. The vertical positioning of the genes is only for ease of plotting and carries no meaning. f) Comparison of the percent of significantly different RNA species in QTL regions and in the genome for the hippocampus. The vertical axis is the percentage of transcripts that are significantly different between the two strains, the horizontal axis lists the cell types. Data from QTL intervals are indicated by dots, colored according to class: excitatory (red), inhibitory (blue) and non-neuronal (grey). Data from the entire genome are indicated by black crosses.

The apparent enrichment of genetically mediated variation in excitatory neurons at the five QTLs containing the causal genes led us to examine whether the same was true for all the QTLs we had identified. We compared gene expression and epigenetic variation at 93 QTLs with the rest of the genome, expecting that any enrichment would be confined to, or more prominent, at the QTLs. Surprisingly, although the QTLs occupy about 0.01% of the genome, the percentage of significantly different transcripts (again defined as exceeding the corrected 5% threshold) in QTL regions was indistinguishable from the percentage of significantly different transcripts in the genome (Figure 6f), a finding replicated for methylation and snATAC-seq data (Table S5). In other words, there was no enrichment in QTLs for genetically mediated variation in excitatory neurons; all regions of the genome contribute to this phenomenon.

Is the enrichment of genetic variants in excitatory neurons significant? It certainly seems so, but at least two confounds could produce this result. For the snATAC-seq and snRNA-seq data, identification of significant strain differences is more likely with higher read coverage, and for methylation the more sites we identify, the higher the likelihood that some will coincide with a sequence variant. We compared a null model, in which sequence coverage predicted the number of differentially expressed genes, ATAC peaks, or methylation sites with mutations, to one that additionally included cell type, and found highly significant improvements in fit (all P-values < 2.2e-16), demonstrating that cell type was an important contributor, over and above the contribution of sequence coverage.

Second, the inclusion of more excitatory cell types than inhibitory could bias findings (eight excitatory cell types compared to three inhibitory in the hippocampus) since the addition of extra cell types increases the chances of finding variant sites. Against this, we note that in the amygdala there were more inhibitory than excitatory cell types (eight versus six). We also ran a test of this hypothesis, by comparing the contribution of class (“Excitatory” and “Inhibitory”) to cell type, considering that cell type was completely nested within class. To do so we resorted to a Bayesian analysis which revealed a significant effect of class (the 95% confidence intervals do not overlap zero). Full results of this analysis, together with a more detailed consideration of the impact of sequence coverage and cell type involvement on the detection of significant strain differences, are presented in Supplemental Material.

Is the enrichment in excitatory cells a peculiarity of the difference between C57BL/6J and DBA/2J? We answered this question for the hippocampus using methylation data from six other strains. Using CAST/EiJ as an outgroup, we compared the number of mutations in CpG sites at excitatory and inhibitory neurons in the ventral hippocampus. Table S6 shows that for each strain compared to CAST/EiJ the proportion of mutated CpG sites was higher in excitatory than inhibitory neurons, supporting our finding that at least for methylation, alterations in functional elements directly attributable to genetic variation were enriched in excitatory neurons compared to other cell types. Together, these results suggest a model whereby genetic effects act on specific hubs of circuits/cell types.

## DISCUSSION

We have established the feasibility of combining mapping with QC testing for gene identification, breaking open a bottleneck in the genetic dissection of complex traits in mice, and provided a set of six genes to further mechanistic understanding of fear-related behavior. Unexpectedly, genetically mediated variation in three independent molecular analyses, methylation, snRNA-seq and snATAC-seq, was found to occur preferentially in excitatory neurons, and to be enriched in the hippocampus compared to the amygdala. These observations demonstrate the possibility of moving quickly from locus to gene, identify novel biology underlying fear-related behavior, and raise questions about expected relationships between genetic and functional variation.

Our findings are important particularly for behavioral research, for which there is a near absence of a physiological understanding of the origins of behavioral variation. At a locus on chromosome 13, we made the unexpected discovery that a lncRNA (*4933413L06Rik*) is involved in cue fear conditioning. Little is known about *4933413L06Rik* (one study found it to be differentially expressed during development of auditory forebrain^69^). Our finding opens new avenues for clarifying the function of this lncRNA in fear-related behavior. Two other causal genes (*Psip1*, *Sh3gl2*) have also never been studied in the context of fear-related behaviors. We note that of the five genes with known functions, four (*Lsamp*, *Ptprd*, *Nptx2*, and *Sh3gl2*) are implicated in synapse development or function^70–74^. Ptprd belongs to the type IIA Receptor-type Protein Tyrosine Phosphatases (LAR-RPTP) family of phosphatases, involved in synaptic formation and structure during development and neurogenesis^71^. Lsamp is a cell surface adhesion molecule that is involved in neuritogenesis and axon guidance, also implicated in the maturation of serotonergic^75^) and thalamocortical circuits^76^. Nptx2 is a secreted glycoprotein that localizes to both pre- and post-synaptic compartments of excitatory synapses containing AMPA receptors, and is involved with synaptogenesis ^77^ and adult neurogenesis^78^. *Sh3gl2* (endophilin1), a cytoplasmic Src homology 3 (SH3) domain-containing protein^79^, localises to presynaptic nerve terminals where it functions in synaptic vesicle endocytosis^80^ and the regulation of exocytosis^81^. The significance of these functional observations is difficult to assess, but it at least suggests that variation in fear-related behavior in part originates in synaptic development.

The QC test could serve as a gold-standard for gene identification following QTL mapping in inbred mice. Following mapping, the construction of knockouts is relatively straightforward, and the test requires a simple breeding protocol. We recommend powering the QC tests based on results from the mapping experiment (the effect size of the locus in the mapping population will be lower than that in QC test^82^) but note that our failure to detect a candidate gene at a locus on chromosome 5 could be due to low power, or because we tested the wrong gene. It is also important to bear in mind that while a positive QC test result reveals a gene through which a QTL has its effect, it does not necessarily mean that the QTL is the one containing the candidate gene. The QC test will detect the effect of any QTL that requires the gene to be present. Typically, that QTL will be the one local to the gene tested, but that assumption is not essential for the functioning of the QC test.

One tantalizing corollary emerging from the analysis of gene causality in fear-related behaviors is that genetic variation may act preferentially in excitatory rather than inhibitory neurons. We had expected that the pattern of strain differences might point to the involvement of a particular cell type, in one region of the brain. Instead of finding cell type enrichment we observed that strain differences in three different modalities, snRNA-seq, methylation and snATAC-seq, were more prevalent in excitatory than inhibitory neurons, particularly in the hippocampus, compared to the amygdala. We found this to be true not just for the QTL regions but for the entire genome. There was little evidence of strain differences in non-neuronal tissues, although we can be less confident in this assertion, because our sample was enriched for neuronal cells and may not be representative of non-neuronal variation.

What might explain this finding? One possibility is that it is specific to a comparison between C57BL/6J and DBA/2J. We think this is unlikely and provide some evidence against this idea by running a similar analysis for methylation data in other strain comparisons, though we have not extended this observation to other modalities. Alternatively, the finding represents a preference for genetic effects in excitatory neurons. Why might this be so?

The idea that genetic effects operate preferentially in excitatory neurons fits with a neuronal circuit model where inhibitory neurons sculpt behaviors driven by excitatory neurons. Excitatory neurons are extremely diverse, resulting in high-dimensional and sparse coding^83^, consistent with the expectation that such an architecture permits highly efficient information transfer^84^. Consequently, encoding within excitatory networks cannot be predicted from their transcriptome alone. By contrast activity within inhibitory neuronal networks is lower dimensional, with correlations between inhibitory cell types determined primarily by cell type, so that cells belonging to a single transcriptomic class are highly correlated^85–87^. In this model, genetic variation is more permissible in excitatory neuronal networks than inhibitory, suggesting that disturbances of inhibitory networks are more likely to be pathological. If this interpretation is correct, identifying genetic effects that disturb function at a circuit level may be more fruitful than looking for cell-type specific genetic effects.

## Acknowledgements

This work was funded in part through NIH grants R01MH115979, R01MH125252, U01MH130995, NIH DP1DA044371 and UCLA Jonsson Comprehensive Cancer Center and Eli and Edythe Broad Center of Regenerative Medicine and Stem Cell Research Ablon Scholars Program. We would like to gratefully acknowledge the UCLA Behavioral Testing Core for assisting with the behavioral experiments, the Cedars-Sinai Applied Genomics, Computation & Translational Core for assisting with the sequencing experiments, and the Genome Editing Core at Augusta University for assisting with the generation of the mouse models necessary to support this research.

## Author contributions

R.C. performed the mouse breeding experiments. C.L. performed the single cell methylation analyses, data from which was processed by M.H. and Z.C. P.C. performed the snRNA and snATAC analyses, with data processing by Z.C. J.F., P.C and D.S. analysed data from the single cell experiments. J.F. and N.L. analysed the mouse mapping data. J.M. provided statistical support. J.F. and P.C. designed the study. P.C., J.F. and E.M. co-wrote the manuscript, which was subsequently reviewed and edited by the rest of the authors.

## Declaration of interests

The authors declare no financial or other competing interests as well as affiliations that are not included in the author list.

## Data availability

- Raw and processed sequencing data generated for this study were deposited to NCBI GEO/SRA and are publicly available at the time of publication. Reviewer links are included here:

`o` https://dataview.ncbi.nlm.nih.gov/object/PRJNA1030354?reviewer=3nknise6o3vtbdfpn385amoaco
`o` https://dataview.ncbi.nlm.nih.gov/object/PRJNA990778?reviewer=b2jb4jjeqvtegl87o3q5u7rue
- Mapping data (including imputed genotypes of all mouse inbred strains and F1s, together with results from the meta-analysis of the three phenotypes)

`o` https://figshare.com/s/7fcb1eebfd9c607afd12
- Knockout interaction data, including phenotypes and genotypes for all quantitative complementation tests

`o` https://figshare.com/s/cc86ed5777cb95b99cec
- Processed single nucleus epigenetic and gene expression data

`o` https://figshare.com/s/39e1c671ec0dc09d5b27

## METHODS

### Mouse phenotyping and mapping

#### Phenotypes

Published data were obtained from three sources: the Jackson Laboratory website (https://phenome.jax.org), the Gene Network website (https://genenetwork.org) and PubMed. Pubmed articles were identified using search terms including ‘fear’, ‘anxiety’, ‘elevated plus maze’, and ‘fear conditioning’. This identified potentially useful data sets from 25 publications ^16–40^, four unpublished data sets at the Jax website and one unpublished data set at the Gene Network website. Processing of each phenotype data set and decisions on which to include are described in **Supplemental Material.** A list of the strains is given in Supp Table 1 and the studies we chose, together with the number of animals and the phenotype names, is given in Table S2.

#### Mouse population

Mice from the Mouse Diversity Panel (HMDP) were used for the behavioral analyses. Mice (n = 700) were obtained through Jackson Laboratory at approximately 60 days old and housed for a 14-day acclimation period prior to testing. Mice were housed in groups (3-4 per cage) under a 12hr/12hr day/night cycle with ad libitum access to food and water. Testing was carried out between 10 AM and 4 PM. Auditory background stimulus in the form of white noise (80db) was delivered through overhead speakers. All protocols conformed to NIH Care and Use Guidelines and were approved by UCLA’s Animal Care and Use Committee (protocol number ARC-2018-026).

#### Behavioral tests

All tests were completed at the Behavioral Testing Core at UCLA. Animals were handled for 5 days prior to experiments. The order of experiments for genetic mapping was EPM -> FC cue/context. EPM -> 7 days-> FC training -> 24 hours -> FC context -> 24 hours -> FC tone.

#### EPM

Animals were placed into an elevated plus maze apparatus (<u>arms 5 cm x 30 cm</u>) for 5 minutes with 15 lux at the center. We measured time spent in open/closed arms, and total entries into open/closed.

#### Fear-conditioning

Mice were placed in the conditioning chamber (context A) for 3 min before the onset of the discrete conditioned stimulus. The parameters were as follows:

Tone: pure tone, 2700Hz, 80dB, rise/fall 50, duration 30s

Shock: 0.75 mAmps, duration 1s, inter-trial interval 180s

Animals were exposed to three pairings of the CS and US. After the CS-US pairings, the mice were left in the conditioning chamber for another 60 s and then placed back in their home cages. FC boxes and software to control the boxes were purchased from MedAssociates.

#### Fear-conditioning context test

24 hours after conditioning, mice were returned to the same context (context A) with no shock or tone and freezing was measured for 8 minutes.

#### Fear-conditioning cued test

24 hours after the context test, mice were placed in a novel context (context B) where, they are exposed to the same procedure as the training but without any shocks (3 minutes exposure followed by 3 CS-US pairings, with parameters equal to fear acquisition). Freezing was measured during the initial 3 minutes exposure, during tone, and during the inter-trial interval.

#### Behavioral analysis

Behavior was recorded digitally from a camera mounted above each test chamber at 30 FPS. Following the completion of experiments, each recorded EPM video was analyzed using ANY-maze for positional tracking of the animal along the apparatus. For EPM mapping we measured open arm entries, time spent in open arms, and closed arm. VideoFreeze (MedAssociates) was used to measure animal freezing in QC testing from recorded videos. Freezing was determined as an absence of all visible movement except that required for respiration; in VideoFreeze, we set the motion threshold to 18 au with a minimum freeze duration of 1 second (30 frames). Contextual fear was assessed by total freezing time over the first 3 minutes of the test. Cued fear was assessed by total freezing time in response to the presentation of the first, second, and third CS during ITI. One exception was with freezing measurements during genetic mapping with BXD lines. Due to the variable coat color across lines, we used ANY-maze for animal tracking and freezing measurements as it performed better than VideoFreeze.

#### Genotypes

We carried out imputation using EMINIM, using the recommended parameter settings for inbred mice given by the software documentation. Roughly 1% of SNPs were triallelic and were discarded. EMINIM returns probabilities for the minor allele, the major allele, and for a missing genotype. In order to perform subsequent GWAS, we translated these results into hard genotype calls. Based on the observation that EMINIM tended to output high confidence values for one of the alleles, we translated a 90% or higher confidence for an allele into a hard call for that allele, with lower confidence calls being called as missing. We combined these imputation results with the genotyped data.

#### Genetic Mapping

All phenotypes were mapped in a two-stage approach. First, we used a mixed model, implemented in GEMMA ^46,47^ (obtained from https://github.com/genetics-statistics/GEMMA), to map each study separately. Input files for GEMMA were generated using PLINK ^88^, obtained from https://zzz.bwh.harvard.edu/plink. We used the PLINK binary format of the required files, including all alleles with a frequency greater than 10%. A genetic relationship matrix was generated using the command:

gemma -bfile input.file -gk -o kinship.file

Mapping was carried out using the command:

gemma -bfile pl input.file -k kinship.file -lmm -o output.file

After mapping with GEMMA, the results files were combined with purpose written perl scripts into a format suitable for meta-analysis with METASOFT ^48,49^. METASOFT (downloaded from http://genetics.cs.ucla.edu/meta_jemdoc/) was invoked with the following command:

java -jar Metasoft.jar -input input.file -mvalue -output output.file

GEMMA assigns effects to the minor allele, as determined by the .bim files from PLINK which means there is considerable variation between studies as to which allele gets an effect (due to allele frequency variation). After running each study though GEMMA we swapped alleles to the most common arrangement, before proceeding to the meta-analysis.

METASOFT calculates a genomic-control inflation factor for the mean effect (lambda_Mean Effect) and a genomic-control inflation factor for heterogeneity (lambda_heterogeneity). These values were calculated for each run, and the software was run a second time, to include the effect of these variables. For example:

java -jar Metasoft.jar -input input.file -mvalue -output output.file -lambda_hetero 2.738124 - lambda_mean 2.430179

#### Significance threshold for the genome-wide association studies

Simulations to estimate the appropriate significance threshold were carried out using GCTA ^50^, obtained from https://yanglab.westlake.edu.cn/software/gcta. Phenotypes were simulated using real genotype data under a simple additive genetic model. A set SNPs was chosen as the causal variants that were seen to have effects in the mapping studies. We obtained estimated effect sizes for these SNPs from GEMMA output files and converted the beta estimate from the linear model to the values required by GCTA using

Beta_gcta = beta_orig * sqrt(2*p*(1-p))

(where p is the allele frequency from the GEMMA output file)

Input SNP files were those used for the mapping with GEMMA. 1000 simulations were carried out using the command:

gcta64 --bfile input.SNP.file --simu-qt --simu-causal-loci causal.snplist --simu-hsq 0.2 --simu-rep 3 --out

After simulating data for each component study, files were processed through GEMMA and METASOFT exactly as for the real data.

To investigate the contribution of genotype structure to the inflation seen in quantile-quantile plots we simulated data in the same way, but with no causal SNPs.

### Generating knockout animals for QC testing

#### Gene selection at loci

We identified genes lying within QTLs, and determine whether they are expressed in brain tissue from in situ hybridization data (Allen Brain Atlas^55^, the mouse ENCODE transcription dataset^56^), and RNA-seq data obtained from the following publications: ^57–62^. For ISH data from the Allen dataset, we looked for presence of expression. For ENCODE transcriptome data we set a cutoff for an RPKM >5 in at least one brain region category, and for RNA-seq studies we ranked expression of each gene and averaged across studies; any gene in the top 25% of all genes (or >rank 6000) was considered expressed. For the Allen institute scRNA-seq dataset, we only included sequencing data from cell type clusters from CA1-CA3 of the hippocampus. Genes that passed expression thresholds for all three datasets were included for further consideration.

#### Mouse KO generation

To generate each KO, we used a CRISPR-Cas9 genome editing approach. For each gene, two single guide RNA (sgRNA) were generated by Synthego Corp. (Redwood City, California, USA) for targeting, except for 4933413L06Rik where we carried out a knock-in with a single-stranded DNA synthesized by IDT, Inc (Coralville, Iowa, USA). Guide RNAs (gRNAs) were designed to maximize both efficiency and specificity scores ^89^. The sgRNA and/or ssDNA, and Cas9 protein (Alt-R® SpCas9 Nuclease V3 from IDT) were co-injected into zygotes of C57BL/6J mice (Jackson Laboratory, Stock#009086). After microinjection, zygotes were transferred into the oviduct of pseudo-pregnant Swiss Outbred mice (Jackson Laboratory, Stock#034608) to generate founder mice. Founder mice were obtained and confirmed by PCR genotyping and Sanger sequencing. The correctly targeted founders were bred with C57BL/6J mice, and PCR genotyping and Sanger sequencing were again performed to confirm germline transmission. Full descriptions of each knockout design, sgRNA sequences, and genotyping primers are given in **Supplemental Material**

To confirm the effect of the deletion on RNA production at each locus we ran quantitative polymerase chain reactions using a standard RT–qPCR assay (Taqpath COVID19 Combo Kit). We analysed RNA extracted from the hippocampus of heterozygous knockouts on C57BL6/J animals.

#### Quantitative complementation testing

We generated offspring from four crosses ^4^. C57BL/6J animals were mated to DBA/2J, and to heterozygote C57BL/6J animals where one allele was the knockout. DBA/2J animals were also mated to the heterozygote knockout mice. This generated four groups: wildtype C57BL/6J, F1 DBA/2J / C57BL/6J (wildtype), heterozygote C57BL/6J knockouts, and F1 DBA/2J / C57BL/6J (knockout). All four groups were assayed for the relevant phenotypes associated with the QTL, using the phenotyping protocols described above.

Evidence that the gene is the QTL gene was then detected as a statistical ‘cross’ (*mutant* or *wildtype*) by ‘strain’ (DBA/2J or C57BL/6J) interaction in a linear model, using the R statistical programing language ^90^. Batch and sex were included as covariates, using the R command

summary (lm (phenotype ∼ sex + batch + cross * strain, data = data))

### Single-nucleus sequencing of ventral hippocampus and amygdala

#### Animals

Male mice from eight inbred strains A/J, C57BL/6J, BALB/cJ, FVB/J, DBA/2J, WSB/EiJ, PWK/PhJ, and CAST/EiJ mice were purchased from Jackson Laboratories at 8 weeks of age and transferred to UCLA where they were kept for at least 7 days before tissue extraction. Animals were housed with *ad libitum* food and water in a 12-hour light-dark cycle.

#### Ventral hippocampus microdissections

Adult male animals (Jackson Laboratories) were euthanized at 10-16 weeks old in an isoflurane chamber and decapitated. The brain was removed, and the ventral region of the hippocampus was micro-dissected, snap frozen in dry ice, and stored at −80 until processing. Tissue from ∼2 animals were combined into a single tube and considered a replicate, with 2 replicates per strain for snmC-seq2, snRNA-seq, and snATAC-seq experiments.

#### Amygdala microdissections

Adult male animals (Jackson Laboratories) were euthanized at 10-16 weeks old in an isoflurane chamber and decapitated. The brain was removed and coronal brain slices containing amygdala tissue were generated on a 1mm brain matrix (World Precision Instruments). Amygdala tissue was micro-dissected from these slices under a dissecting scope in cold PBS, snap frozen in dry ice, and stored at −80 until processing. Tissue from ∼2-3 animals were combined into a single tube and considered a replicate, with 2 replicates per strain for snmC-seq2, snRNA-seq, and snATAC-seq experiments.

#### Generating snmC-seq2 libraries

We carried out snmC-seq2 on micro-dissected tissue as previously described ^65^. Briefly, frozen tissue was homogenized into single nuclei suspensions with Dounce homogenization, then immediately sorted on into a 384-well plate with a FACSAria sorter (BD Biosciences) at the UCLA Flow Cytometry Core. We selected for a 75-25 enrichment of neuronal vs non-neuronal nuclei during FACS sorting using NeuN-488/DAPI counterstains (Millipore Sigma MAB377X). Bisulfite conversion and single cell methylome libraries were generated following this step.

#### Generating snRNA-seq libraries

Single nuclei suspension and library generation were completed at the Cedars Sinai Applied Genomics, Computation and Translational Core and followed the 10X protocol for the Chromium Next GEM Automated Single Cell 3’ Library and Gel Bead Kit v3.1 (cat# PN-100014) as described except for the following modifications:

Suspensions from cell nuclei were generated using the recommended method from the 10X scMultiome protocol (CG000375 Rev C) to lyse cells and obtain nuclei. Following single nuclei suspension generation, nuclei were counterstained for 7-AAD and NeuN-405 antibody (Novus Biologicals, 1:200) and sorted on a MACSQuant Tyto (Miltenyi Biotech) prior to GEM generation. We selected for a 75-25 split of NeuN+/7-AAD+ nuclei for neurons and NeuN-/7-AAD+ for non-neuronal nuclei respectively.

We captured ∼10,000 nuclei per genotype per region per replicate on a single 10X GEM chip. All downstream library preparation was done according to the 10X Genomics protocol (CG000286) and sequenced on a Novaseq 6000 with a target of ∼40-50k reads per nucleus.

#### Generating snATAC-seq libraries

Single nuclei suspension and library generation were completed at the Cedars Sinai AGCT core and followed the 10X protocol for Next GEM scATAC-Seq v1.1 (PN-1000175) as described except for the following modifications:

Nuclei suspensions were generated using the recommended method from the 10X scMultiome protocol (CG000375 Rev C) to lyse cells and obtain nuclei.

Following single nuclei suspension generation, nuclei were counterstained for 7-AAD and sorted on a MACSQuant Tyto prior to GEM generation. NeuN was not used for neuronal enrichment due to dye incompatibility between our NeuN antibody and a nuclear counterstain. After the sort, we carried out permeabilization of nuclei as per the protocol. We aimed to capture 10,000 nuclei per well x 8 wells, for a total of 80,000 nuclei over 8 total samples (∼10,000 nuclei per genotype per region per replicate). All downstream library preparation was done according to the 10X Genomics protocol (CG000209) and sequenced on a Novaseq 6000 with a target of >35k reads per nucleus.

### Analysis of single-nucleus datasets

#### Mapping and primary quality control

All reads were mapped to the mouse mm10, Genome Reference Consortium Mouse Build 38 (GCA_000001635.2)). The gene and transcript annotation used was a GENCODE GTF file ^91^. snmC-seq2 reads were mapped using Bismark (V0.22.3) to SNP-swapped mm10 genomes. This was done to allow us to directly compare chromosomal coordinates between the 8 strains. SNPs for each strain were downloaded from the Sanger Institute website^41^. Custom code was used to generate 8 separate SNP-swapped genomes for each strain by replacing each corresponding SNP nucleotide position in the B6J mm10.fa file with the nucleotide of that strain. snRNA and snATAC-seq reads were mapped using 10X Cell Ranger (V6.0.2) and 10X Cell Ranger ATAC (V2.0.0) respectively against mm10 for B6 and a SNP-swapped mm10 for DBA. We retained introns for RNA analysis while default settings were used for ATAC analysis.

#### Single-nucleus RNA sequencing data quality control and pre-processing

All quality control and pre-processing were done under the Seurat package framework ^92^. Per biological sample, we filtered out cells that (1) fall below the 5th percentile of the total UMI counts (nCount_RNA) or the 5th percentile total number of unique genes expressed (nFeature_RNA) or 700 unique genes expressed, whichever was more stringent; (2) are over the 95th percentile quantile in terms of either the total UMI counts or the total number of unique genes expressed; (3) have larger than 5% mitochondria fraction (percent.mt).

Global coverage normalization: counts per million (CPM) was applied to each cell followed by log transformation (“LogNormalize”). We then projected cells from each biological sample to low dimensional space using principal components analysis (PCA) on highly variable features selected by Seurat. Potential doublets were identified and subsequently removed from the downstream analysis by DoubletFinder ^93^, ran in the top 15 principal components space with the expected doublet rate set to the recommended amount from 10X genomics based on high quality yield volume.

#### Single-nucleus ATAC sequencing data quality control and pre-processing

All quality control and pre-processing were done under the Seurat and Signac package framework ^94^. Per sample, we first used Model-based Analysis for ChIP-Seq (MACS) to call sample specific de-novo peaks from its fragments file^95^. We then merged sample-specific sets of peaks to a unified peaks set while removing peaks with length larger than 10000bp or smaller than 20bp. A unified peaks by cells count matrix was constructed from the fragments file while removing cells with lower than 200 peaks detected and peaks only present in less than 10 cells. Cells were filtered based on the following criteria: (1) appropriate number of non-duplicate, usable read-pairs (passed_filters from Cell Ranger’s output singlecell.csv). Specifically, we set it to larger than 3000, 4000, 2500, and 5000 for the 2 BL6 and 2 DBA samples collected in the hippocampus region. Similarly, we set it to be larger than 7500, 5000, 7500, and 7500 for their amygdala counterpart. (2) number of fragments overlapping peaks (peak_region_fragments from Cell Ranger’s output singlecell.csv) falls within the 5th percentile and the 95th percentile. (3) ratio of fragments overlapping peaks over the total number of non-duplicate, usable read-pairs falls within the 5th percentile and the 95th percentile. (4) nucleosome_signal: the ratio of fragments between 147 bp and 294 bp (mononucleosome) to fragments < 147 bp (nucleosome-free) is smaller than 4 (5) TSS enrichment score is larger than 2. We did not include a filter for ratio of peaks in black list regions over the total number of non-duplicate, usable read-pairs as this was removed during the construction of the DBA SNP-swapped reference genome. We normalized the count data with Text Frequency Inverse Document Frequency (RunTFIDF) and performed Singular Value decomposition (RunSVD) on top 90% informative features selected by Signac (FindTopFeatures). The first low dimensional embedding was excluded from downstream doublet detection and clustering analysis due to high correlation with sequencing depth. Potential doublets were identified and subsequently removed from downstream analysis by DoubletFinder ^93^, ran on the 2^nd^ - 11^th^ low dimensional embedding with the expected doublet rate set to the recommended amount from 10X genomics based on high quality yield volume. Finally, we built a gene-by-cell transcriptional activity matrix that counts per cell, at the gene body and 2000bp upstream to capture the promoter region, the total number of ATAC-seq counts.

#### Single-nucleus methylation data quality control and pre-processing

Cells were filtered on the basis of several metadata metrics: (1) mCCC level <0.03; (2) global mCG level >0.5; (3) global mCH level <0.2; (4) total mapped reads >100,000; (5) Bismarck mapping rate >0.5; and 6 (percent genome covered > 2). Methylation features were calculated as fractions of methylcytosine over total cytosine across gene bodies ± 2kb flanking regions and 100kb bins spanning the entire genome. Methylation features were further split into CG and CH methylation types. Features overlapping our methylation mm10 blacklist were removed. 100kb bin features were then filtered on minimum mean coverage >500 and maximum mean coverage <3000. Gene body features were filtered on minimum coverage >5 and all remaining features were normalized per cell using the beta binomial normalization technique in allcools ^67^. Individual CpG sites were only counted when they had ≥5 reads covering the site.

#### Single-nucleus RNA sequencing data integration, clustering and annotation

Gene counts were normalized using SCTransform ^96^, and regressed out percentage of reads from mitochondrial genes. We then integrated cells from all samples using reciprocal principal components analysis (rPCA) implemented in Seurat 4.0.5 ^92^ on the top 5000 integration genes and in the top 30 reciprocal principal components space. For clustering, we standardized the integrated data, performed PCA on all integrated genes and ran de-novo Louvain clustering algorithm in the top 15 principal components space with resolution set to 0.1. Cluster markers that are conserved between the strains were called using non-parametric Wilcoxon rank sum test and subsequently used for annotation. We annotated hippocampals clusters by manually checking conserved markers against the ALLEN BRAIN MAP’s Mouse Whole Cortex and Hippocampus dataset. For cells collected in the amygdala, we performed a second round of analysis restricted to those identified to be neuronal in the initial annotation to increase our annotation resolution. We used the same integration pipeline and parameters described above except when clustering we ran *de-novo* Louvain clustering algorithm in the top 25 principal components space with resolution set to 0.75, due to the increased expected complexity of cell types in this brain region. All clusters were first grouped into broad types of inhibitory or excitatory neurons except for one cluster that simultaneously expressed Gad2 and Slc17a7. We thus simply annotated it as “Neuronal” followed by its by cluster marker gene names. We also manually compared our clusters to those that were previously reported in amygdala ^97^, and adapted their annotation for clusters with high concordance in top marker genes.

#### Single-nucleus ATAC sequencing data integration, clustering and annotation

We first jointly projected all cells’ ATAC peak profile to uncorrected Latent Semantic Indexing (LSI) embeddings with TFIDF transformation followed by calculating SVD on the top 90% most informative peaks. Peak profile embeddings were then integrated in shared low dimensional space via integration anchors identified in the 2^nd^ to 30^th^ reciprocal LSI space as implemented in Signac 1.5.0. We then integrated cell transcriptional activity profiles by performing SCTransform after regressing out percentage of activity from mitochondrial genes and carried out rPCA integration on integration genes identified from the single-nucleus RNA experiment and in the top 10 reciprocal principal components. We transferred the single-nucleus RNA annotation onto the single-nucleus ATAC cells by linking the RNA’s expression profile with ATAC’s transcriptional activity profile through canonical correlation analysis (CCA) described in Seurat. Pairs of cells from each modality that are mutual nearest neighbors in the top 15 canonical components space were identified as “transfer anchors”. “Anchors” were further filtered and weighted by distances in the integrated peak embeddings prior to imputing ATAC cells’ annotation of cell type. For amygdala cells, after the initial assignment to cell type, we repeated the above-mentioned pipeline only on those cells predicted to be neuronal to obtain a set of higher resolution annotation. The second round of analysis was performed on integration genes identified from the subset of single-nucleus RNA data also categorized as neuronal.

#### Single-cell methylation data integration, clustering and annotation

We used the negative of the average mCH fraction of the gene body ± 2kb as the proxy of methylation cells’ transcriptional activity as described previously ^67^. We first integrated, across strains, single-nucleus methylation gene-body mCH profiles on RNA integration genes via “integration anchors” identified in the top 30 reciprocal PCA space. Gene body mCH profiles were then linked to single-nucleus RNA expression profiles via “transfer anchors” identified in the top 15 canonical components space. “Transfer anchors” were further filtered and weighted by distances in the integrated mCH embeddings prior to imputing methylation cells’ annotation using their RNA counterpart as reference. For amygdala, we directly linked to the higher resolution annotation used on the neuronal cells in RNA. These served as our final annotation for all the neuronal cells in both hippocampus and amygdala. Since mCH methylation, used to construct the “transfer anchors”, is largely not present in the non-neuronal population^98^, we further performed de-novo annotation on methylation cells using both the genome wide mCG and mCH fraction at 100kb as described previously^99^. Non-neuronal clusters were annotated by their canonical marker genes (gene-body hypo-methylation). These served as our final annotation for all the non-neuronal cells in both hippocampus and amygdala.

#### Co-embedding of single-nucleus RNA, single-nucleus ATAC, and single-nucleus methylation sequencing data

Both single-nucleus methylation and single-nucleus ATAC cells’ expression profiles were imputed with previously computed “transfer anchors”. All three modalities were merged on their integrated or imputed expression profiles, projected to low dimensional space via PCA, and visualized by UMAP (performed on the top 15 principal components).

#### Cell-type specific differential test for single-nucleus ATAC and single-nucleus RNA

We used DESeq2 ^68^ for cell-type specifc pseudobulk level differential expression analysis and differential accessibility analysis. Per cell type, raw counts were aggregated to replicate level and DESeq2 was run under default parameters to detect statistical evidence of strain differences. Any genes with no coverage in either or both strains were excluded.

#### Testing class and cell type effects on strain differences

We used the *brms* package in R to test for the effect of class analysis^100^. We defined a Bayesian model using *brms* with a negative binomial distribution for the number of significant differences (for RNA and ATAC data) or the number of mutations for methylation. We used up to eight chains and visually inspected the output to ensure chains were mixed. We used the following code to define the model:

brm(significant_difference ∼ 1 + (1 | class/celltype) + log(coverage), data = data, chain = 8, control = list(adapt_delta = 0.95), family = negbinomial(link = “log”))

## Selection of data sets for the meta-analysis

We describe here the pipeline we used to harmonize the phenotypic data. Identifying phenotypes for the meta-analysis had to deal with the heterogeneity of the data sets collected. Each experiment used different protocols, different testing equipment and often reported different, though related, assays. Different studies use different names for what is supposedly the same phenotype. For example, for fear conditioning some studies reported the time freezing^1^, other reported its opposite, the amount of activity^2^. Some record 3 minutes of freezing, some five minutes, or more. EPM behavior is recorded in apparatus of different dimensions, under different lighting conditions, at different times of day, all of which are known to affect behavior ^3^. We dealt with this heterogeneity as follows.

After downloading data, and examining what was most commonly collected, we decided to use the time freezing during a five-minute period after exposure to an altered context (contextual freezing), and the mean time spent freezing during exposure to a conditioned stimulus (most usually a tone). For the elevated plus maze we used the number of entries into the open arms. We created a data dictionary to convert the different phenotype names to a common set, and this is provided in Table S2.

Some studies also provided information about sex and age, and we tested, using a linear model, whether there was an effect on phenotypes of either. Since we found no significant effects (at a 5% threshold after correcting for multiple testing) and since some studies did not provide this information, we did not include covariates in downstream analyses.

Our most challenging problem was to determine whether phenotypes that we labelled as the same, really were the same. We examined the relationship between phenotypes from different studies in two ways. First, for each strain for each study we obtained the trait mean and then calculated the correlation between the same strains in different studies. Not every study tests the same strains, so although there were 42 possible pairwise comparisons for FC-context, only 30 comparisons could be carried out. One study (Palmer5) was dropped because of insufficient overlap in the strains used with other studies.

There was a considerable range in the correlations between studies (from −0.04 to 0.93), though only 13 comparisons were significant (at a P <0.05, uncorrected for multiple testing), in part reflecting differences in the variation in the number of strains in each comparison (from 3 to 31). Table 1 summarizes the findings and demonstrates that in some cases a pair of studies are correlated, but not with other studies (for example for the CUE phenotype in the table, study Bolivar2 is correlated with Bothe-2005 (r = 0.85) but neither study is highly correlated with others).

**Table 1:** phenotypic correlations between strains in different studies for three phenotypes: Cue fear conditioning (CUE), contextual fear conditioning (CONTEXT) and entries into the open arms of the elevated plus maze (EPM). Significant (P <0.05) results are shown in bold type.

| CUE | Brigman-2009 | Owen-1997 | Bolivar2 | Yang-2008 | Smith-2011 | Bothe-2005 | Knoll-2016 | Carhuatanta-2014 | UCLA |
| --- | --- | --- | --- | --- | --- | --- | --- | --- | --- |
| Brigman-2009 | 1 | 0.04 | NA | 0.67 | 0.45 | NA | **0.70** | 0.27 | 0.08 |
| Owen-1997 |  | 1 | NA | 0.23 | 0.21 | NA | 0.31 | 0.42 | 0.36 |
| Bolivar2 |  |  | 1 | -0.02 | 0.15 | **0.85** | NA | NA | 0.16 |
| Yang-2008 |  |  |  | 1 | 0.27 | NA | 0.92 | **0.50** | 0.10 |
| Smith-2011 |  |  |  |  | 1 | 0.14 | 0.11 | 0.30 | 0.53 |
| Bothe-2005 |  |  |  |  |  | 1 | NA | NA | 0.21 |
| Knoll-2016 |  |  |  |  |  |  | 1 | **0.68** | 0.30 |
| Carhuatanta-2014 |  |  |  |  |  |  |  | 1 | 0.37 |
| UCLA |  |  |  |  |  |  |  |  | 1 |

| CONTEXT | Bolivar2 | Smith-2011 | Carhuatanta-2014 | Yang-2008 | Brigman-2009 | Bothe-2005 | Knoll-2016 | Owen-1997 | UCLA |
| --- | --- | --- | --- | --- | --- | --- | --- | --- | --- |
| Bolivar2 | 1 | 0.45 | NA | NA | NA | **0.93** | NA | NA | 0.13 |
| Smith-2011 |  | 1 | 0.07 | **0.64** | 0.09 | 0.72 | 0.18 | 0.21 | 0.14 |
| Carhuatanta-2014 |  |  | 1 | 0.43 | 0.30 | NA | 0.49 | **0.54** | **0.56** |
| Yang-2008 |  |  |  | 1 | 0.65 | NA | 0.34 | 0.29 | 0.40 |
| Brigman-2009 |  |  |  |  | 1 | NA | **0.46** | 0.52 | **0.35** |
| Bothe-2005 |  |  |  |  |  | 1 | NA | NA | 0.17 |
| Knoll-2016 |  |  |  |  |  |  | 1 | 0.25 | 0.16 |
| Owen-1997 |  |  |  |  |  |  |  | 1 | 0.46 |
| UCLA |  |  |  |  |  |  |  |  | 1 |

| EPM | Brown<br>1 | Flaherty<br>1 | Wahlsten1 | Schalkwyk<br>1 | Shin1 | Milner-2008 | UCLA |
| --- | --- | --- | --- | --- | --- | --- | --- |
| Brown1 | 1 | -0.04 | 0.14 | 0.27 | 0.24 | 0.05 | 0.24 |
| Flaherty1 |  | 1 | 0.50 | 0.27 | 0.70 | 0.44 | 0.76 |
| Wahlsten1 |  |  | 1 | **0.73** | 0.01 | **0.85** | 0.33 |
| Schalkwyk1 |  |  |  | 1 | 0.23 | 0.61 | -0.09 |
| Shin1 |  |  |  |  | 1 | 0.11 | 0.73 |
| Milner-2008 |  |  |  |  |  | 1 | **0.69** |
| UCLA |  |  |  |  |  |  | 1 |

As a second measure of the relationship between studies, we ordered the means of each strain, and ranked them. We then applied the order for one study to another to examine the extent to which studies agreed on the order of strain means. We decided to use the pattern of correlations to select studies for the genetic analysis, requiring the presence of more than 5 strains in common, and a positive correlation greater than 0.2 (though we did not require this to be significant since). The studies we chose, together with the number of animals and the phenotype names, is given in Supp Table 2.

## Imputation

This section describes the results of the imputation method we used to fill in missing genotypes in the inbred strains used for mapping. The highest density set of genotyped markers is that derived from 16 strains for which we have complete sequence data ^4^ but this does not include any recombinant inbreds and lacks some of the strains used for mapping. Since the missing strains are descendants of a small number of founder haplotypes ^5^, we can impute the genotypes in the non-sequenced strains. Using genotypes from arrays as a scaffold ^5–7^ we imputed missing genotypes using data from sequenced animals. Most imputation algorithms are designed for fully outbred populations, where a key problem is correctly phasing haplotypes (e.g. ^8^). Alternative methods are needed for efficient imputation in mouse strains and so we used the software tool EMINIM ^9^, which requires a reference file, containing strains with known haplotypes, and a target file for imputation, containing strains genotyped at some SNPs. We prepared these two files by placing strains that were genotyped at all SNPs in the reference file, while strains with one or more missing SNPs were placed in the target file. For the BXD mice, SNPs between genotype markers from the same founder haplotype were pre-filled due to the negligible likelihood of multiple recombination events between consecutive genotyped SNPs. This resulted in 27 reference strains and 227 target strains. We then ran EMINIM imputation on each chromosome separately and obtained calls on 16,767,664 SNPs (triallelic SNPs, constituting 1% of the total, were discarded). Since EMINIM outputs high confidence values for one of the alleles, we translated a 90% or higher confidence for an allele into a hard call for that allele, with lower confidence calls being called as missing. We combined these imputation results with the genotyped data. The rate of missing calls was less than 0.8% for 18 of 19 autosomal chromosomes (chromosome 19 had a missing call rate of 1.03%).

We validated the imputation results by comparing imputed calls to genotypes from sequenced BXD strains obtained from ^10^. The sequence-derived set contained 4,325,552 SNPs, of which 3,812,095 variant sites overlapped with our imputation. Across all chromosomes, each BXD strain was imputed with between 93% and 99% accuracy. Imputation accuracy for all strains combined varied between 89% and 99% per chromosome, with 96.29% overall. Due to the pre-filling procedure described previously, this accuracy reflects only the most challenging SNPs to impute: those between genotyped SNPs from different founder haplotypes. When including the genotyped and pre-filled SNPs, the accuracy for each chromosome was greater than 99.64%.

We performed a second assessment of imputation accuracy by removing the C57BL/6J and DBA/2J strains from our reference panel, randomly masking 50% of the bases of those strains, and using the remaining inbred strains in the reference panel to impute the masked bases. Once again, we used EMINIM with its default inbred imputation parameter settings and translated a 90% or higher confidence call for an allele into a hard genotype call. Calls with less than 90% confidence were translated to a missing call, leading to a missing genotype rate of 0.09% to 0.59% per chromosome. The accuracy of imputation on the masked bases was 95.6-99.7% per chromosome, and 97.9% overall.

## Design of CRISPR-Cas9 knockouts

In this section we describe the design of the knockouts used for the quantitative complementation experiments and include the primers used to detect them along with evidence from polymerase chain reaction amplification for their detection. We used a CRISPR-Cas9 genome editing approach to delete the genes, as described in the Methods section. Here we give details of the guide RNAs (sgRNA), and genotyping primers, showing the results for each gene. Where an existing knockout had been successfully created, usually on a C57BL/6N background, we adopted the same design, targeting the same part of the gene but in the C57BL/6J strain. We genotyped each animal to confirm deletion of the targeted region and used quantitative PCR to confirm that the deletion affected abundance of RNA transcript in hippocampal tissue from F1 animals, as described in Methods.

### 1) Hcn1 KO design - Delete exon 2

We targeted Hcn1 using the strategy described in^11^ to generate the Hcn1tm1a(KOMP)Wtsi. sgRNA sequences:

sg769r: 5’- TCAATTTAGCGATCACAGAA *AGG* −3’

sg1673r: 5’- ACACACTCAAATAGTCGTCA *TGG* −3’

Genotyping primers:

Hcn1 wtF1: 5’-GGTCCAGCATCTAGTCCAG-3’ Hcn1 wtR1: 5’-GCAGCAACGAGGTCTGACT-3’

Wt = 2.145 kb; mt ∼ 1.2 kb

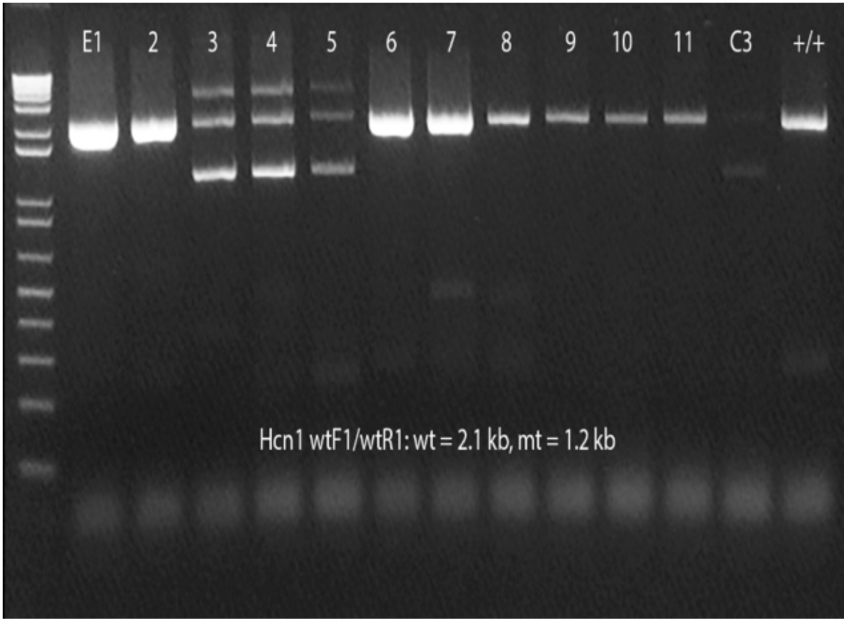

### 2) Megf9 KO design - Delete exon 2

We used the same approach as in ^11^, Megf9tm1a(KOMP)Wtsi, to design our Megf9 deletion. sgRNA sequences:

sg352r: 5’- CGTATTCTAAGCCAATGTAG TGG −3’

sg940r: 5’- TTAAATAACTGATTCTCCAC AGG −3’

Genotyping primers:

Megf9 wtF: 5’- GCCATTTCCAGCATGAGCTCCAAG −3’

Megf9 wtR: 5’- GGTAGAAGCAGAGATCTGACTCCTGT −3’

Wt = 1077 bp; mt ∼490 bp

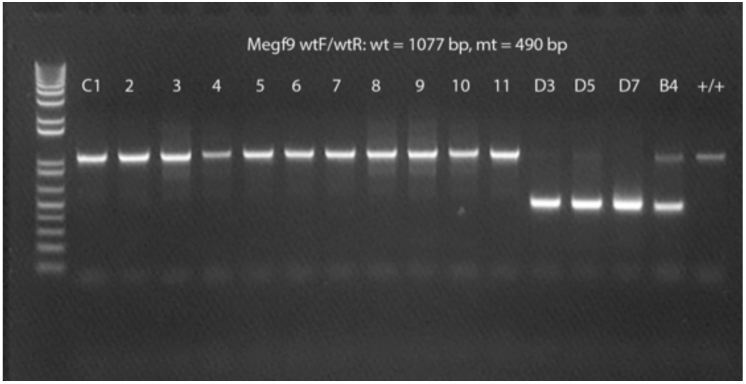

### 3) Ptprd KO design - Delete exon 22

We used the same approach as in ^11^, Ptprdtm2a(KOMP)Wtsi, to delete Ptprd, where these KO animals have been used in ^12^, where RNA expression was used to confirm deletion. sgRNA sequences:

sg559: 5’- GGCAGAATGTATTCTTAATC TGG −3’

sg960r: 5’- GTTAAAAAGTGCTACCCTGG GGG −3’

Genotyping primers:

Ptprd wtF2: 5’-GAGGGTTGGGCACTGAGAAG-3’ Ptprd wtR1: 5’-GCCTGGTTAGACCTTGCTGA-3’

Wt = 764 bp; mt ∼360 bp;

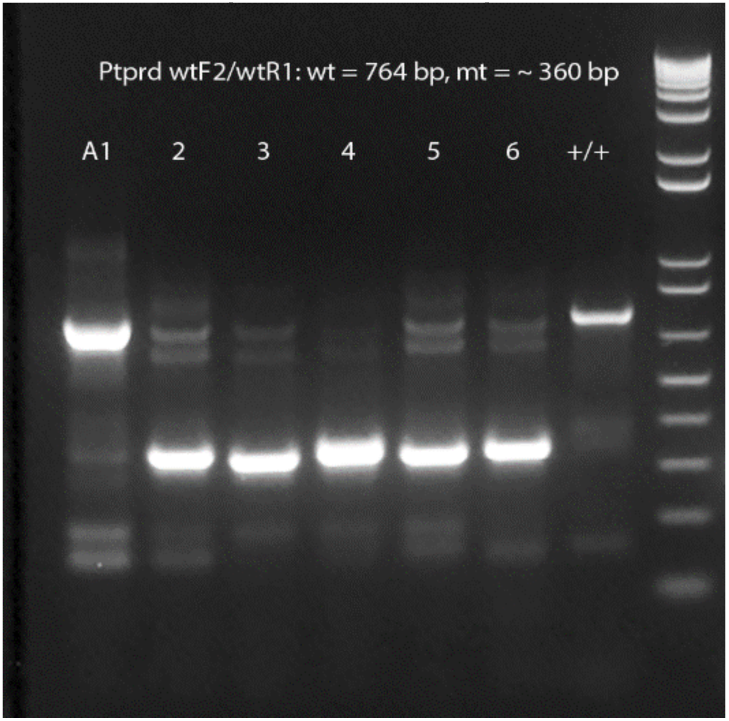

### 4) Stim2 KO design - Delete exons 6-7

We used a similar approach as in ^13^, which targeted exons 4-7 of Stim2 for deletion, where gene deletion was confirmed by Western blot. We deleted exons 6-7.

sgRNA sequences:

Stim2 sg370r: 5’- AAACCACATATGGCTATAAC AGG −3’

Stim2 sg1620: 5’- GGAGCAGGTACCGTCCTAGT GGG −3’

Genotyping primers:

Stim2 wtF: 5’- CTGATTCTCCAGCTTGTGCCTAGTGACA −3’

Stim2 wtR: 5’- GCCGAGGAACAGCAGACATGACA −3’

Wt = 1475 bp, mt= 234 bp

Stim2 wtF2: 5’- GTGACAATGACCTGGCCTAACTGAAG 3’-

Stim2 wtR2: 5’- CCCACCTCCCAGCGTTAGC −3’ (in intron 6-7)

572 bp for wt only

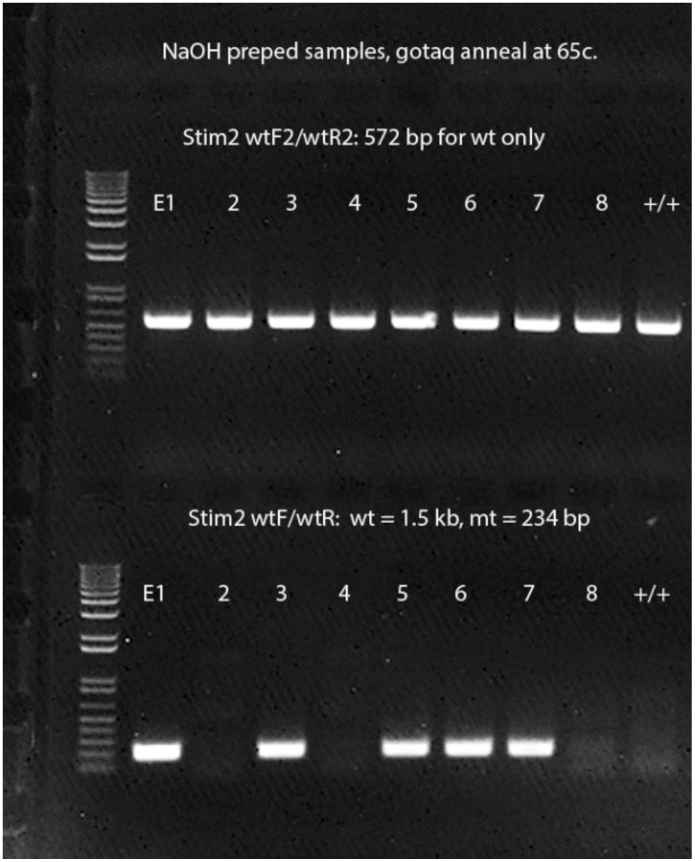

### 5) Nptx2 KO design - Delete exon 1

We used the same approach as in ^14^ to generate knockouts of Nptx2 by targeting exon 1, where gene deletion was confirmed by Northern blot

sgRNA sequences:

Nptx2 sg416r: 5’- CGTTCGCGTCACGCGGAGGG CGG −3’

Nptx2 sg1216: 5’- AAATGGGTACGTGGGCTGGC CGG −3’

Genotyping primers:

Nptx2 wtF1: 5’- GGCAAGTGGAGTCATCTCGGTCAGGA −3’

Nptx2 wtR1: 5’- CCTCGGTAAGTTCACGGTGAGTCTAGTG −3’

Wt = 1242, mt = 451

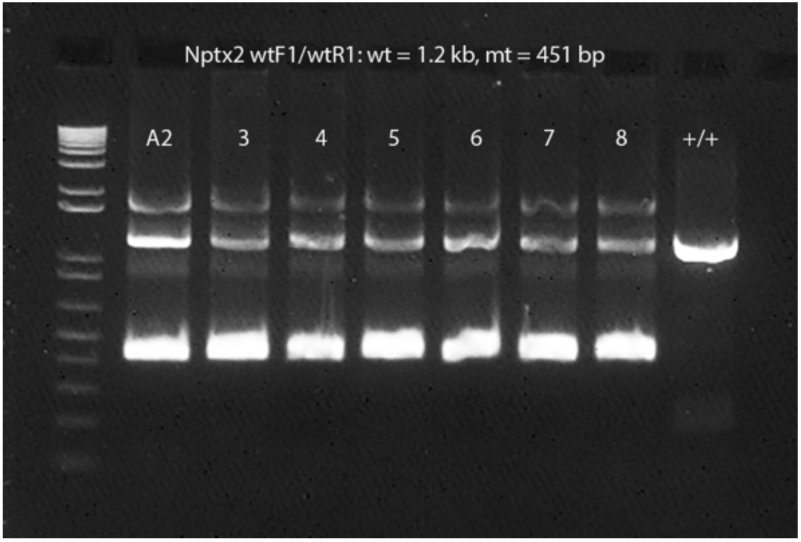

### 6) Lsamp KO design - Delete exon 2

We used the same approach as in ^15^ and^16^ to generate knockouts of Lsamp by targeting exon 2, where gene deletion was confirmed by Western blot. sgRNA sequences:

Lsamp sg433r: 5’- GATTTCATTCTCCGCGTGAC AGG −3’

Lsamp sg793: 5’- CGGCTGTGACGAGTCAAGAG TGG −3’

Genotyping primers:

Lsamp wtF: 5’- CTGAGCCCTCTCAGAAATGTCACGACA −3’

Lsamp wtR: 5’- GTATCACCAGCACATGGAGTTTCCTAGT −3’

Wt = 543 bp, mt = 192 bp

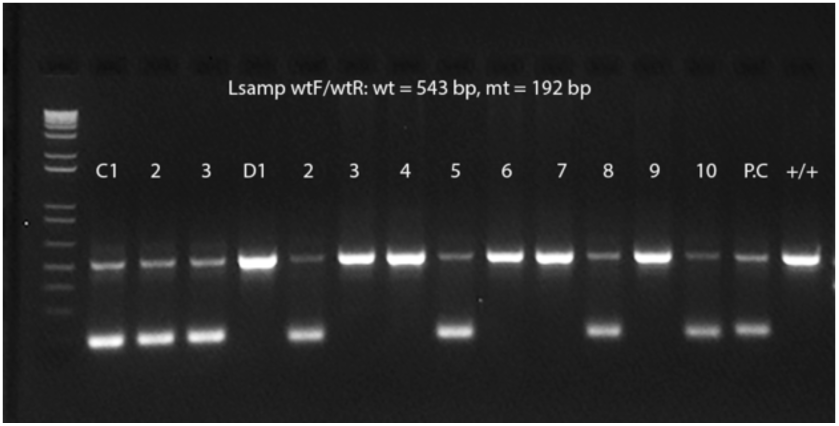

### 7) Snapc3 KO design - Delete exon 3

No known knockout allele was previously reported; we selected exon 3 for deletion. Snapc3 sg560: 5’- ATAGACGCTCACAACTCTGG AGG −3’ (ordered 8/20/21) Snapc3 sg1129: 5’- CGGCTCCCAGGGCATGTCAT AGG −3’ (ordered 8/20/21)

Genotyping primers:

Snapc3 wtF: 5’- GCTCAGTAGTAGAGTGCTTGCCTAGCA −3’

Snapc3 wtR: 5’- CCAGCACTCAGGAGATAGACAGGTGACT −3’

WT = 1,011bp; MT = 442 bp, 65-68c

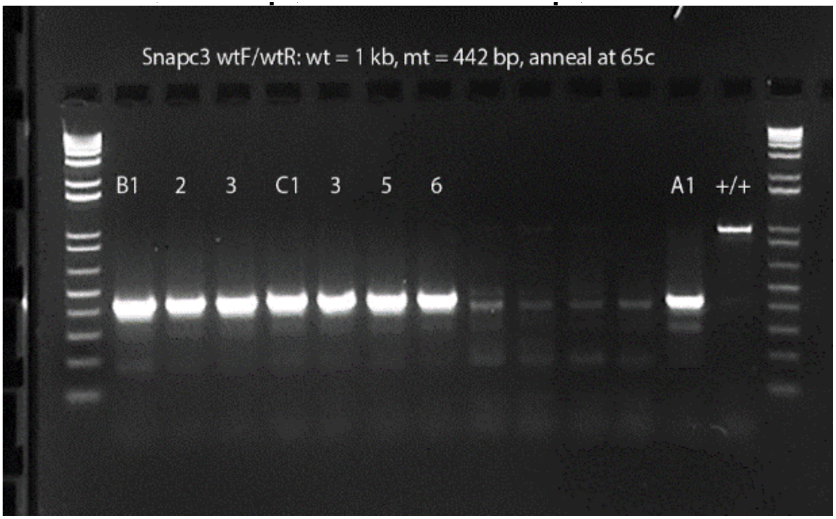

### 8) Sh3gl2 KO design - Delete exon 3

We used the same approach as in ^17^ to generate a knockout by deleting exon 3, which showed loss of expression in brain tissue by Western blot.

sgRNA sequences:

Sh3gl2 sg883r: 5’- AGGCTTCAGGAGCAGTTAGG AGG −3’

Sh3gl2 sg1229r: 5’- ATGGTTAAATACCCCTCAGT TGG −3’

Genotyping primers:

Sh3gl2 wtF: 5-CTGAGGTCTTCAGGCTGGCTGA-3’

Sh3gl2 wtR: 3’-GGCAGTCCTCTATGCCCTATGCA-3’

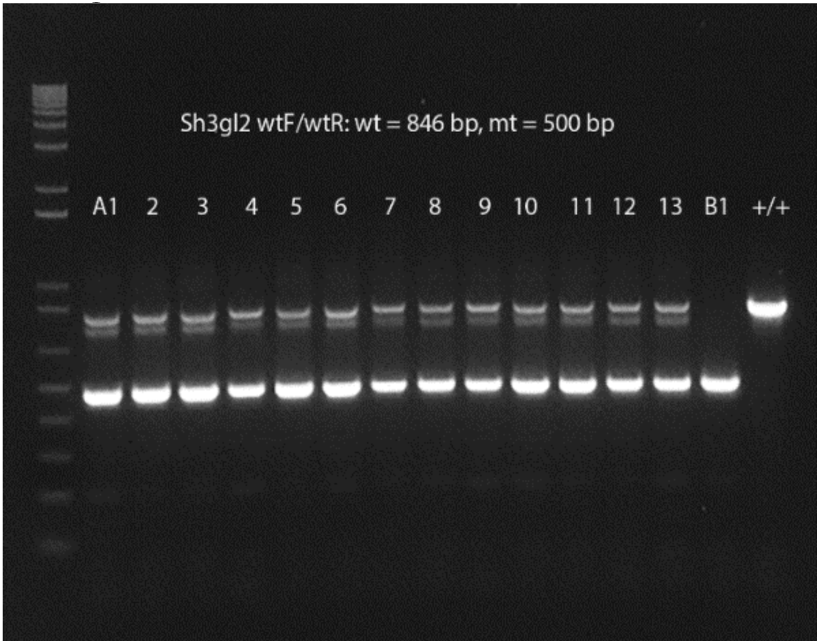

### 9) Psip1 KO design - Delete exon 3

We used the same approach as in ^11^, Psip1tm1a(EUCOMM)Wtsi, to generate our knockout. sgRNA sequences:

Psip1 sg864: 5’- TTTCAATGGAGCTGATCTGG GGG −3’

Psip1 sg1245r: 5’- TCAGGCTAGCCGGGTCTCAT TGG −3’

Genotyping primers:

Psip1 wtF: 5’-GGTCTTGATGTCAGGACCAGT-3’

Psip1 wtR: 5’-CCATGAGCTCAGGCTAGCTCA-3’

WT = 761 bp; MT = 377 bp 62-64c

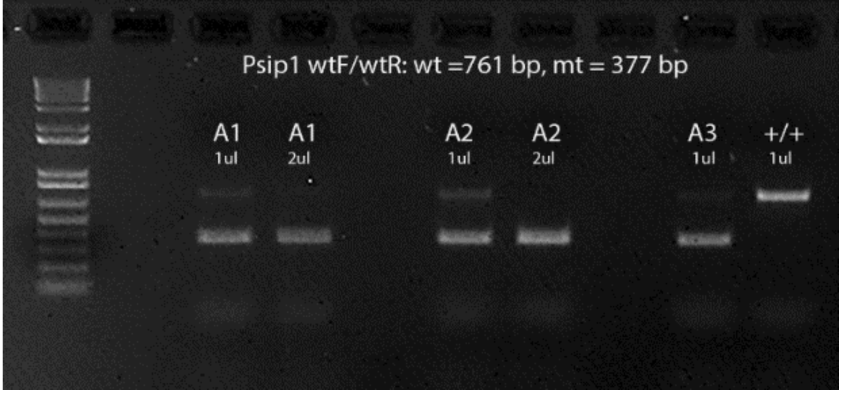

### 10) Ttc39b KO design - Delete Exon 2

We used the same approach as ^11^, Ttc39btm1e(EUCOMM)Wtsi, in generating our knockout. sgRNA sequences:

Ttc39b sg888r: 5’- GCGGGCAGTAACAATCCTAG TGG −3’

Ttc39b sg1299: 5’- TCTCAGTGTTTTGAGCGTAG TGG −3’

Genotyping primers:

Ttc39b wtF: 5’-GACTGATGTGGGCATGTCCTGCT-3’

Ttc39b wtR: 5’-CCTGTGCCACGACCTCCTAGCT-3’

WT = 942 bp; MT = 534 bp

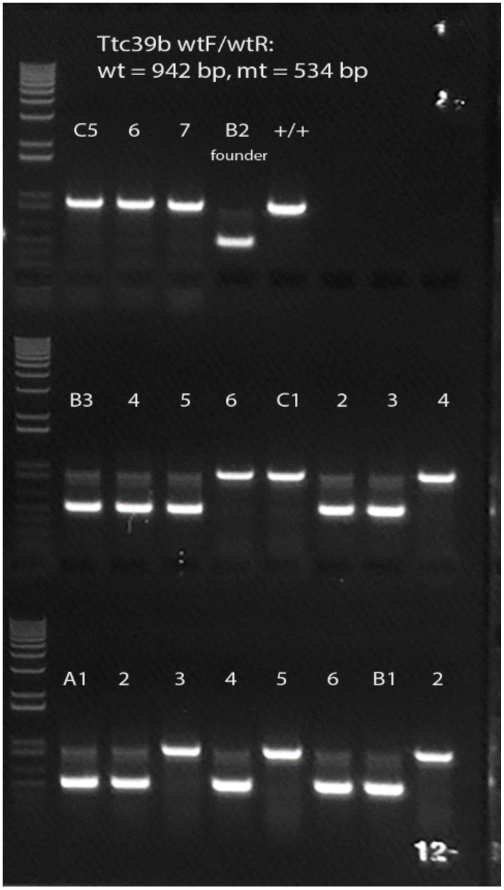

### 11) 4933413L06Rik lncRNA KO through polyA insertion in exon 1

We inserted an SV40 polyA sequence (cloned from Addgene pGP-AAV-syn-jGCaMP8s-WPRE) into exon 1 of 4933413L06Rik using an ssODN-mediated knock-in approach ^18^.

sgRNA sequences:

4933413L06Rik sg1092: 5’- AAAATGTGGTCATTTCCACA TGG −3’

4933413L06Rik KO ssODN (SV40 pA sequence in italics):

5’-CGACAGAAAAATGGATACAGAAAATGTGGTCATTTCC*taagatacattgatgagtttggacaaaccaca actagaatgcagtgaaaaaaatgctttatttgtgaaatttgtgatgctattgctttatttgtaaccattataagctgcaataaacaagtt*A CATGGAATACTACTTGGCTATTAAGAATGAGGACATCCT −3’

Genotyping primers: (ordered 9/30/21)

4933413L06Rik F: 5’- GTCACAAGGGCACATGTTCCACT −3’

4933413L06Rik R: 5’- CCCCCACTGCTCACCAC −3’

Wt = 434 bp, mt = 556 bp

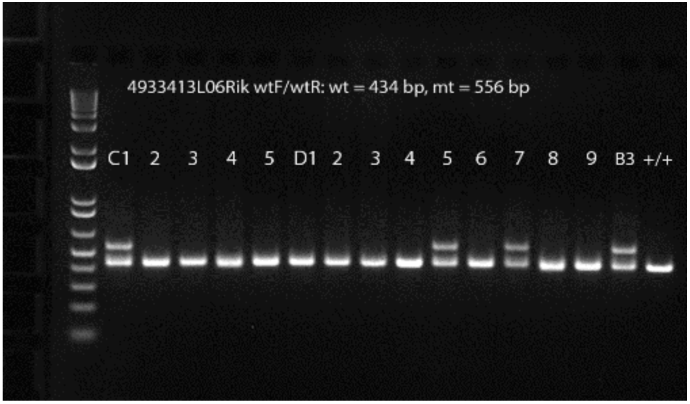

### 12) Parp8 KO design - Delete Exons 20-23

We used the same approach as ^11^, Parp8<tm1.1(KOMP)Wtsi>, in generating our knockout. sgRNA sequences:

Parp8 sg925r: 5’- CTTGCATGATAATCGGCAAA AGG −3’

Parp8 sg3511: 5’- AGGTAACCTGAAAGCACGTG TGG −3’

Genotyping primers:

Parp8 wtF: 5’-GAGCCACTGTCAACTCTGTCAGCTCA-3’

Parp8 wtR: 5’-GCCAGCAAACAGGAGGTCATTGTCTCCA-3’

wt = 3,007 bp; mt = 426 bp

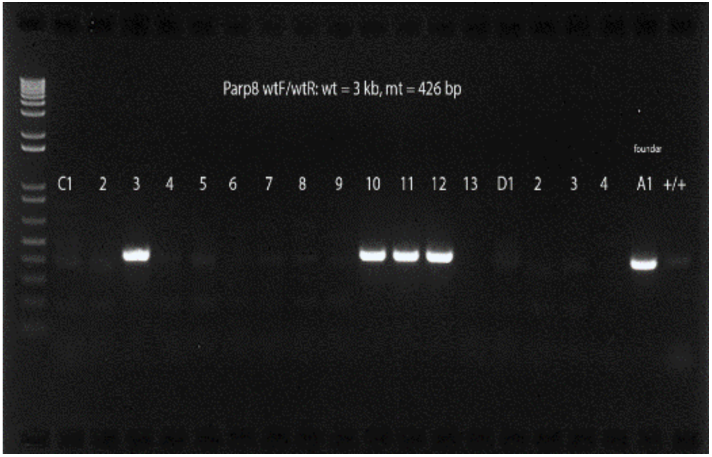

### 13) Mrps30 KO design - Delete Exon 2

No known knockout allele was previously reported; we selected exon 2 for deletion. sgRNA sequences:

Mrps30 sg1055: 5’- TCTCATCTTATTCCGGTGAG TGG −3’

Mrps30 sg1492: 5’- CCGGTGTGAGCATTGCCAAA AGG −3’

Genotyping primers:

Mrps30 wtF: 5’-GGTAGTACAGAAGGTTGAACTCAGGGAC-3’

Mrps30 wtR: 5’-GCTCACCGGCACTAACATCTGCTGATG-3’

WT = 1,063 bp; MT = 626 bp

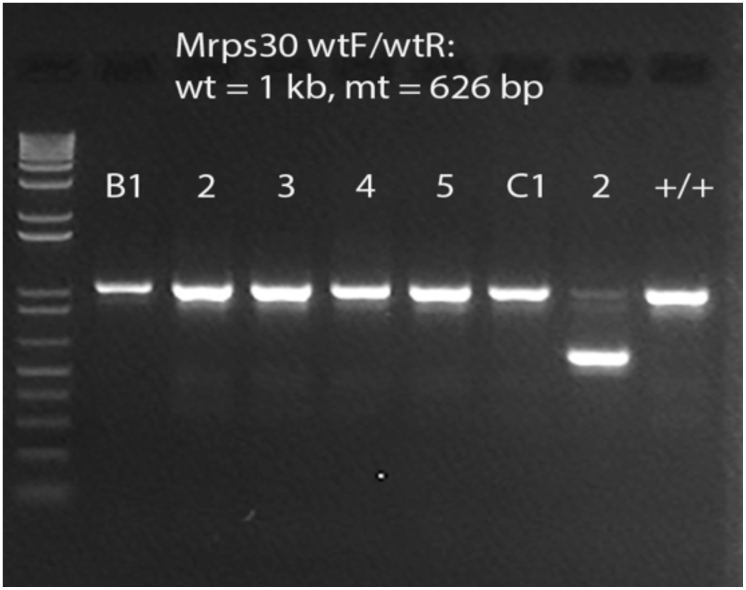

### 14) Emb KO design - Delete Exon 5

We used the same approach as ^11^, Embtm1a(KOMP)Wtsi, to generate our knockout. sgRNA sequences:

Emb sg775r: 5’- ATCATCTGCCGGTGTACAAC TGG −3’

Emb sg1375: 5’- CAAGTGGATTGAAGTTTCGA AGG −3’

Genotyping primers:

Emb wtF: 5’-CCCTCCACATCGTGCCTCATCCTTCTG-3’

Emb wtR: 5‘-GGTGTACCTGTCTGTAAGGGTGTGAGCA-3’

Wt = 971 bp, mt = 380 bp

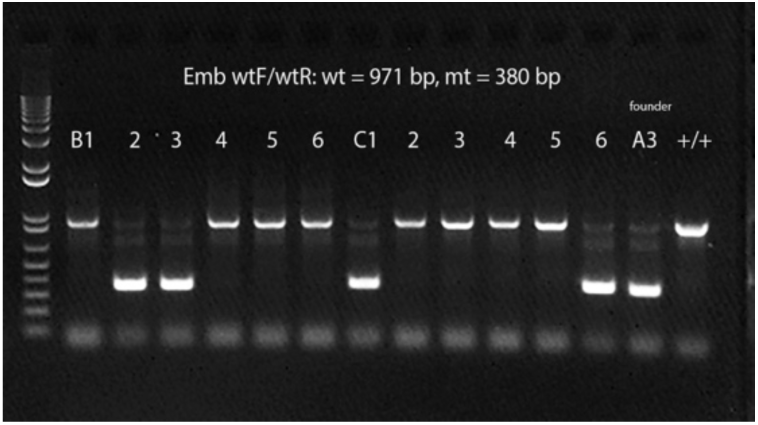

### Genotyping of mouse KO colonies

For genotyping each KO line, we chose different primers that generated shorter products to simplify the genotyping protocol:

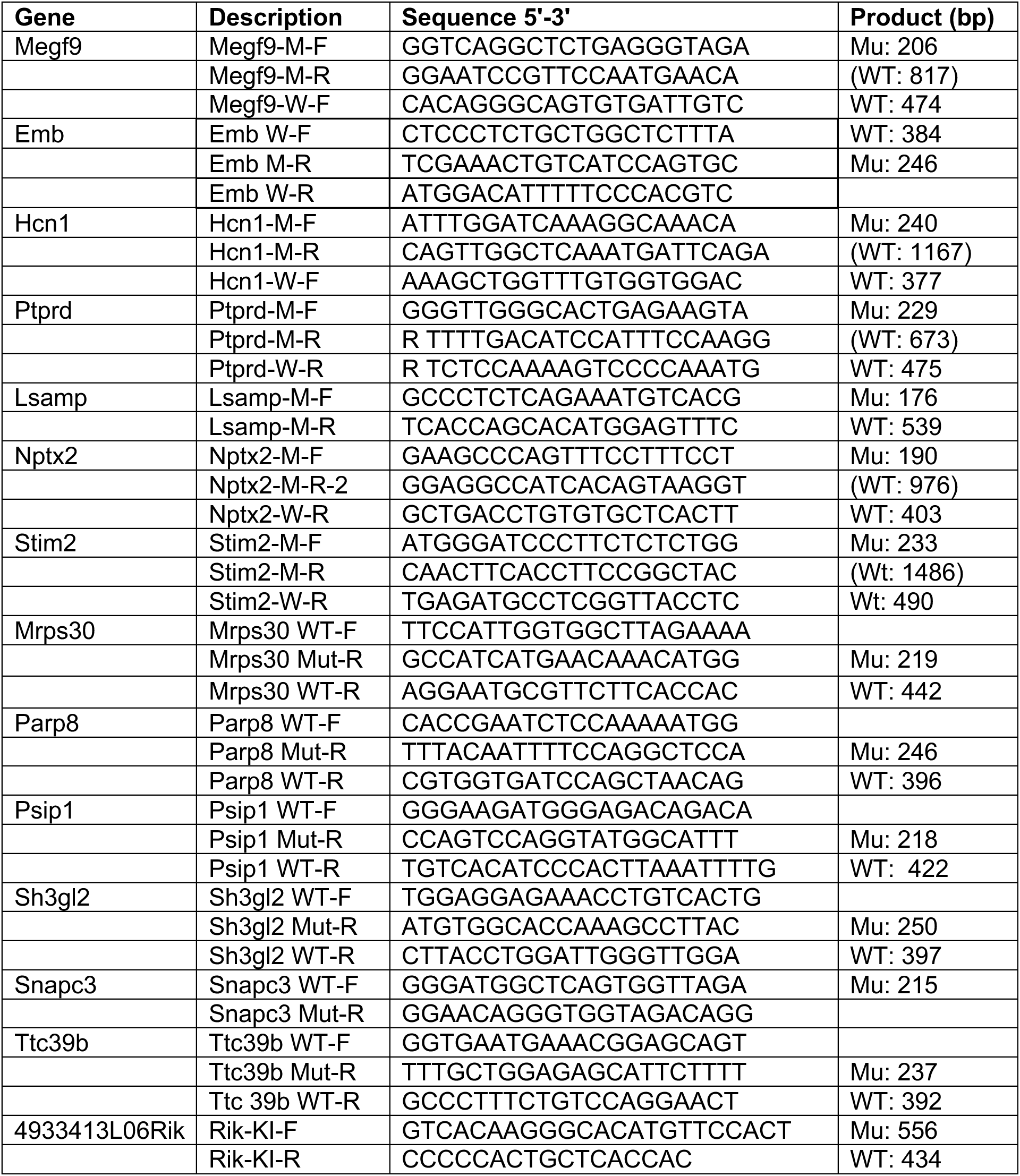

## Enrichment of genetically mediated variation in excitatory neurons

In the main text we report that genetic variation in three epigenetic assays (single nucleus RNA, Assays of Transposase Accessible Chromatin (ATAC-seq) and single nucleus methylation) is more prevalent in excitatory than inhibitory neurons. Here we explore this observation in more detail.

We began by counting the number of RNA species and ATAC sites that differed significantly between C57BL/6J and DBA/2J, and the number of methylation sites containing mutations that removed the methylated site. Significant differences were determined by exceeding an adjusted *p* value below a 10% FDR cutoff value, the default from the output of DESeq2 ^19^). Data for each cell type is shown below for the two brain regions (hippocampus and amygdala), with counts of the significant species, and percentages of the total.

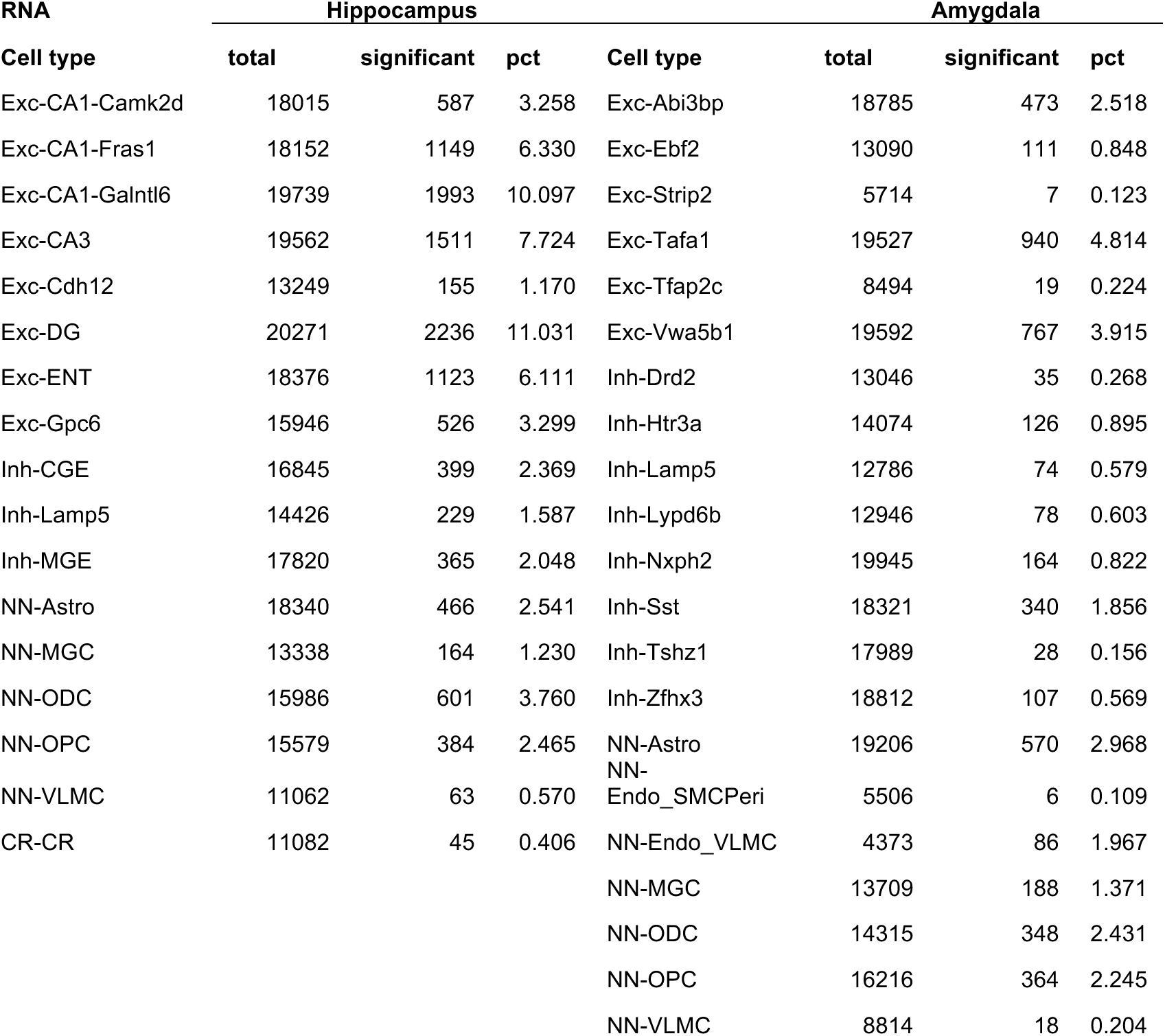
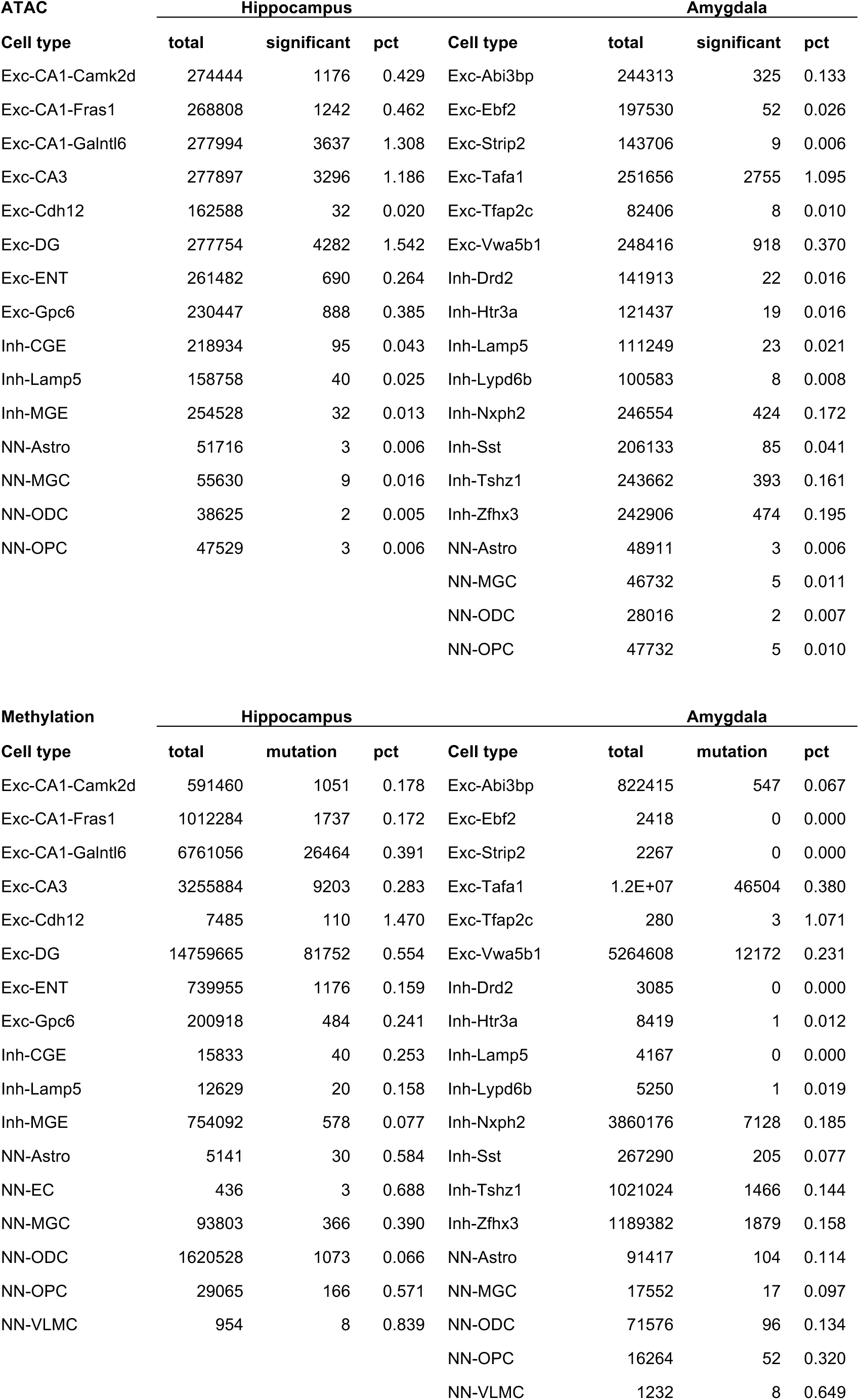

We simplify the above results by taking means of the excitatory and inhibitory cell types from the above data, weighted by the number of observations, to show that there appear to be more excitatory than inhibitory variable species. The table below shows the weighted means of significant strain differences, expressed as a percentage, for the excitatory and inhibitory classes of cell type in the hippocampus and amygdala.

| Modality | Class | Hippocampus | Amygdala |
| --- | --- | --- | --- |
| RNA | Excitatory | 6.47 | 2.72 |
|  | Inhibitory | 2.02 | 0.74 |
| ATAC | Excitatory | 0.75 | 0.35 |
|  | Inhibitory | 0.03 | 0.10 |
| Methylation | Excitatory | 0.45 | 0.32 |
|  | Inhibitory | 0.08 | 0.17 |

Prima facie there is more variation in the excitatory than inhibitory neurons in all modalities, a finding supported by highly significant Fisher’s exact test P-values (shown below with 95% confidence intervals), all of which were P < 2e-16. However, the exact test fails to consider the contribution of differences in sequence coverage, and the presence of more excitatory than inhibitory cell types (8 versus 3 in the hippocampus).

We tested the relative contribution of sequence coverage and cell type by comparing the fit of two models. In one model we used a single predictor, sequence coverage, for the number of significant species. In the second model we added cell type as a predictor. We then tested the improvement in fit using ANOVA in the following way:

fit0 <-glm(number_significant ∼ seq_coverage, data = data, family = binomial(link = “logit”))

fit1 <-glm(number_significant ∼ seq_coverage + celltype, data = data, family = binomial(link = “logit”))

anova(fit0, fit1, test = “Chisq”)

We found in each case the improvement was highly significant, as shown below:

| Modality | Tissue | Deviance | P-value |
| --- | --- | --- | --- |
| RNA | Hippocampus | 1291.6 | <2.2e-16 |
|  | Amygdala | 1282.2 | <2.2e-16 |
| ATAC | Hippocampus | 12895.3 | <2.2e-16 |
|  | Amygdala | 8771.2 | <2.2e-16 |
| Methyl | Hippocampus | 28295.7 | <2.2e-16 |
|  | Amygdala | 517.86 | <2.2e-16 |

While this result indicates that cell type contributes to the number of significant differences independently of coverage, it does not alone support the view that there are more significant differences in excitatory cell types.

Using a generalized linear model (the glm function in R), we tested whether sequence coverage and class (“Inhibitory” or “Excitatory”) predict whether RNA or ATAC sites were significant, and whether methylation sites were mutant or not (in each case coding outcomes as 1 or 0). The results are shown below. The effect of class is relative to “Excitatory” and the direction for each modality is the same (negative, meaning that prediction is positive for “Excitatory”), and highly significant.

| Modality | Tissue | Predictor | Estimate | z-value | P-value |
| --- | --- | --- | --- | --- | --- |
| RNA | Hippocampus | Sequence coverage | 6.15E-05 | 30.80 | <2e-16 |
|  |  | Class | -8.27E-01 | -28.11 | <2e-16 |
|  | Amygdala | Sequence coverage | 6.72E-06 | 5.10 | 3.48E-07 |
|  |  | Class | -3.99E-01 | -13.46 | <2e-16 |
| ATAC | Hippocampus | Sequence coverage | 3.64E-04 | 30.91 | <2e-16 |
|  |  | Class | -2.20E+00 | -50.03 | <2e-16 |
|  | Amygdala | Sequence coverage | 1.65E-04 | 6.92 | 4.53E-12 |
|  |  | Class | -9.51E-01 | -35.44 | <2e-16 |
| Methyl | Hippocampus | Sequence coverage | 1.08E-01 | 102.61 | <2e-16 |
|  |  | Class | -9.06E-01 | -31.96 | <2e-16 |
|  | Amygdala | Sequence coverage | 8.39E-02 | 56.54 | <2e-16 |
|  |  | Class | -6.29E-01 | -61.35 | <2e-16 |

We also ran the generalized linear model using a negative binomial distribution and again obtained highly significant results.

We next tested whether the P-values were well calibrated. We created an empirical distribution by randomly selecting the same number of transcripts as in the observed data for each cell type from the total set of transcripts in all cell types, while ensuring that a gene was selected only once for each cell type. We analysed the empirical data in the same way as the real data, using the generalized linear model to test whether sequence coverage and class (“Inhibitory” or “Excitatory”) predicted the strain differences, and confirmed that the P-values we obtained for the real data were significantly different from the empirical null distribution (P < 0.0001 in all cases).

It’s possible that the enrichment of variable sites in excitatory neurons could be because we identified more excitatory than inhibitory cell types (8 versus 3 in the hippocampus). The presence of more excitatory cell types could increase the chance of revealing additional variable species. This is unlikely to be true since we find the same enrichment in the amygdala tissue, where there are six excitatory neuronal cell types and eight inhibitory, but our analysis has not excluded the possibility of such heterogeneity. To test whether the number of strain differences are significantly different between the two classes (Excitatory and Inhibitory) while considering the coverage difference and accounting for the different numbers of cell types in each class, we used a Bayesian framework.

We used the *brms* package in the software language R for this analysis^20^. We defined a Bayesian model using *brms* with a negative binomial distribution for the number of significant differences (for RNA and ATAC data) or the number of mutations (for methylation data). The model includes predictors for *Coverage* and *Class*. We fitted the Bayesian model using the *brm* function. Results, given in the table below, show that the estimate for class for each tissue and each modality has confidence intervals that do not overlap zero.

| Modality | Tissue | Level | Predictor | Estimate | Est. Error | Lower 95% CI | Upper 95% CI |
| --- | --- | --- | --- | --- | --- | --- | --- |
| RNA | HIPPOCAMPUS | Group | Class | 0.68 | 0.56 | 0.02 | 2.09 |
|  |  |  | Class:celltype | 0.52 | 0.15 | 0.32 | 0.89 |
|  |  | Population | Coverage | 0.50 | 0.01 | 0.49 | 0.51 |
| RNA | AMYGDALA | Group | Class | 0.85 | 0.79 | 0.02 | 2.85 |
|  |  |  | Class:celltype | 1.10 | 0.25 | 0.73 | 1.68 |
|  |  | Population | Coverage | 0.38 | 0.01 | 0.36 | 0.39 |
| ATAC | HIPPOCAMPUS | Group | Class | 2.54 | 1.31 | 0.73 | 5.81 |
|  |  |  | Class:celltype | 1.26 | 0.41 | 0.69 | 2.27 |
|  |  | Population | Coverage | 0.60 | 0.02 | 0.56 | 0.63 |
| ATAC | AMYGDALA | Group | Class | 1.33 | 1.45 | 0.04 | 5.19 |
|  |  |  | Class:celltype | 0.42 | 0.32 | 0.02 | 1.23 |
|  |  | Population | Coverage | 0.70 | 0.14 | 0.44 | 0.97 |
| Methylation | HIPPOCAMPUS | Group | Class | 1.37 | 1.37 | 0.04 | 4.87 |
|  |  |  | Class:celltype | 0.95 | 0.47 | 0.36 | 2.13 |
|  |  | Population | Coverage | 2.12 | 0.31 | 1.52 | 2.74 |
| Methylation | AMYGDALA | Group | Class | 0.86 | 0.79 | 0.04 | 3.07 |
|  |  |  | Class:celltype | 0.49 | 0.33 | 0.13 | 1.39 |
|  |  | Population | Coverage | 2.45 | 0.44 | 1.62 | 3.30 |

Table of posterior samples obtained from the *brms* package. Results are given for the two tissues (hippocampus and amygdala) and three modalities (RNA, methylation and ATAC), reporting standard deviations for group-level effects (class and cell type nested within class) and estimates for population-level effects (sequence coverage). The two-sided 95% credible intervals (lower and upper 95% confidence interval (CI) based on quantiles are given. Results are from eight chains, each of 2,000 iterations of which the first 1,000 are warmup to calibrate the NUTS sampler. To reduce divergent transitions, `adapt_delta` was set to 0.95. Chain convergence was 1, as estimated from the potential scale reduction factor on split chains ( $R_{\text{hat}} = 1$ ).

## Notes

### Competing Interest Statement

The authors have declared no competing interest.

